# Evolutionary rates are correlated between *Buchnera* endosymbionts and mitochondrial genomes of their aphid hosts

**DOI:** 10.1101/2020.11.15.383851

**Authors:** Daej A. Arab, Nathan Lo

## Abstract

The evolution of bacterial endosymbiont genomes is strongly influenced by host-driven selection. Factors affecting host genome evolution will potentially affect endosymbiont genomes in similar ways. One potential outcome is correlations in molecular rates between the genomes of the symbiotic partners. Recently, we presented the first evidence of such a correlation between the mitochondrial genomes of cockroaches and the genome of their endosymbiont (*Blattabacterium cuenoti*). Here we investigate whether similar patterns are found in additional host-symbiont partners. We use partial genome data from multiple strains of bacterial endosymbionts, *B. aphidicola* and *Sulcia mulleri*, and the mitochondrial genomes of their sap-feeding insect hosts. Both endosymbionts show phylogenetic congruence with the mitochondria of their hosts, a result that is expected due to their identical mode of inheritance. We compared root-to-tip distances and branch lengths of phylogenetically independent species pairs. Both analyses show a highly significant correlation of molecular rates between genomes of *Buchnera* and mitochondrial genomes of their hosts. A similar correlation was detected between *Sulcia* and their hosts, but was not statistically significant. Our results indicate that evolutionary rate correlations between hosts and long-term symbionts may be a widespread phenomenon.

## Introduction

The rate of molecular evolution is a fundamental biological trait and is governed by a multitude of factors (Bromham 2009) (Bromham and Penny 2003; Ho and Lo 2013). For example, the substitution rate is influenced by variations in selection and population size, while the mutation rate is influenced by generation and DNA repair mechanisms (Bromham 2009). In symbiotic organisms, evolutionary rates may be influenced by the biology of both host and symbiont. This influence is especially pronounced in strictly vertically transmitted obligate intracellular symbionts (hereafter ‘endosymbionts’), which have a highly intimate relationship with their hosts (Douglas 2010). For example, reductions in the efficacy of selection due to small host population size will potentially lead to increased fixation of deleterious mutations in both host and endosymbiont genomes. One additional factor that could influence rates of molecular evolution in endosymbionts is the dependence of these organisms on the DNA replication and repair machinery of the host, due to loss of the genes encoding these functions in the genome of the endosymbiont (Mao and Bennett 2020). The use of the same DNA replication and repair machinery by host and symbiont could lead to a correlation between the evolutionary rates of the two genomes.

In long-term associations between insect hosts and their endosymbionts, strict maternal vertical transmission and lack of horizontal transfer of symbionts leads to phylogenetic congruence between host and endosymbiont. This provides an opportunity to test for correlations in evolutionary rates between the genomes of host and endosymbiont. To this end, we recently analysed branch lengths on phylogenetic trees for 55 cockroach species and their bacterial endosymbiont *Blattabacterium cuenoti* (Arab et al. 2020). We found evidence of a correlation in evolutionary rates between host mitochondrial and endosymbiont DNA. Although this was the first time such a correlation was reported, previous studies have found correlations in rates between nuclear DNA and either mitochondrial or chloroplast DNA (Hua et al. 2012; Lourenço et al. 2013; Sheldon et al. 2000; Sloan et al. 2012; Smith and Lee 2010; Yan et al. 2019). Since endosymbionts are associated with an array of insect groups, a question arises, is the correlation in evolutionary rates between host mitochondrial and endosymbiont DNA specific to cockroaches? Or is it a more general phenomenon?

Insects that feed on plant sap are among the most diverse groups in the order Hemiptera. Diversification of this group has been attributed to the acquisition of bacterial endosymbionts in multiple lineages. *Buchnera aphidicola* (hereafter ‘*Buchnera*’) is an intercellular bacterial endosymbiont that has been in an obligatory intercellular and mutualistic relationship with aphids for over 200 million years (Douglas 1998; Moran et al. 2008; Moran et al. 1993). These endosymbionts can be found within the body cavity of aphids in highly specialized cells (bacteriocytes) and are vertically transmitted to offspring (Baumann et al. 1995; Braendle et al. 2003; Koga et al. 2012). *Buchnera* is found in almost all aphids, except in cases where it has been replaced with a fungal (Fukatsu and Ishikawa 1996) or another bacterial endosymbiont (Chong and Moran 2018). Genome analyses of *Buchnera* have shown that they range in size from 412 to 645 kb, and have confirmed the important role of these bacteria in provisioning essential amino acids that are lacking in phloem sap, which allowed aphids to tap into these specialized diets (Baumann 2005; Moran et al. 2008). Many host-transcribed genes responsible for amino acid biosynthesis pathways, nitrogen recycling, and transport of nutrients were found to be overexpressed in the bacteriocytes of aphids, indicating a high level of cooperation and integration of host and endosymbiont genomes (Hansen and Moran 2011; Nakabachi et al. 2005).

*Sulcia mulleri* (hereafter ‘*Sulcia*’) is another endosymbiont found in a number of sap-feeding insects in the hemipteran suborder Auchenorrhyncha, including a number of species of cicada, planthopper, treehopper, and spittlebug. Acquisition of *Sulcia* in this group is believed to have occurred ~260 million years ago, making it among the most ancient examples of bacterial symbiosis (McCutcheon and Moran 2007; Moran et al. 2008; Moran et al. 2005). The genome of *Sulcia* ranges in size from 191 to 277 kb, and its gene inventory indicates that *Sulcia* can provision eight of the ten essential amino acids lacking in the host’s xylem or phloem sap diets (Bennett and Moran 2013; McCutcheon and Moran 2010). For that reason, all Auchenorrhyncha infected with *Sulcia* have co-primary bacterial or fungal endosymbionts that are capable of provisioning the remaining two amino acids (Bennett and Moran 2013; Mao and Bennett 2020). A transcriptomic study investigating the bacteriocytes of the glassy-winged sharpshooter and the aster leafhopper found that host genomes appear to provide support to *Sulcia* through enhanced gene expression, including those encoding enzymes involved in replication, transcription, and translation (Mao and Bennett 2020).

*Buchnera* and *Sulcia* are among the best studied insect endosymbionts, with genome sequences available for multiple taxa of each genus. The availability of such data, combined with the availability of mitochondrial genomes for their host taxa, make them ideal candidates for further investigations into whether evolutionary rates among hosts and endosymbionts are correlated. Here we infer the phylogenies of 31 *Buchnera* strains and 28 *Sulcia* strains and compare them with the mitochondrial phylogenies of their hosts. We then compare branch lengths and the rates of evolution for host-endosymbiont pairs across these phylogenies. We find a correlation in evolutionary rates between *Buchnera* and aphids and some evidence that evolutionary rates in *Sulcia* and their hosts might be correlated.

## Methods

### Genomic data from aphids and Buchnera

We obtained 31 annotated whole *Buchnera* genomes, 28 annotated whole *Sulcia* genomes, 28 whole and partial *Sulcia-*host mitochondrial genomes and 15 whole and partial aphid mitochondrial genomes directly from GenBank submissions (table S1). Sixteen aphid whole and partial mitochondrial genomes were assembled from Illumina Hiseq 4000 paired reads obtained from the Sequence Read Archive (SRA) (see table S1). We filtered reads using a local library of 15 published aphid mitochondrial genomes as reference sequences. *De novo* mitochondrial genome assembly was then performed for each sequence library using Velvet v1.2.10 (Zerbino 2010). All mitochondrial genes were annotated using the MITOS web server (Bernt et al. 2013). Across 31 *Buchnera* genomes and 28 *Sulcia* genomes, we identified orthologues using OMA v1.1.2 (Altenhoff et al. 2019).

Using a custom script, we found 240 and 120 protein-coding genes to be shared by all *Buchnera* and *Sulcia* taxa, respectively. For aphid mitochondrial gene alignments, all taxa were represented by at least 10 of 12 available protein-coding genes. For *Sulcia-*hosts, missing genes were presumed to be a result of the relatively low sequencing coverage used, leading to uneven sequencing coverage of samples. Genome sequences of outgroups for *Buchnera* alignments were obtained from GenBank and included three strains of *Escherichia coli* (NC_000913.3), one *Serratia marcescens* (HG326223.1) and one *Yersinia pestis* (NC_003143.1). Genome sequences of outgroups for *Sulcia* alignments were obtained from GenBank and included *Flavobacterium gilvum* (CP017479), *Lutibacter* sp. (CP017478), *Tenacibaculum dicentrarchi* (CP013671). The *Daktulosphaira vitifoliae* (DQ021446.1) mitochondrial genome was used as an outgroup for aphid alignments, while *Trialeurodes vaporariorum* (AY521265) was used as an outgroup for *Sulcia*-host alignments.

TranslatorX (Abascal et al. 2010) was used to align the 240 and 120 orthologous genes at the amino acid level individually and concatenated into 249,624 bp and 128,214 bp alignments, for the *Buchnera* and *Sulcia* alignments, respectively. The mitochondrial genome data set included all protein-coding genes from each taxon plus the genes encoding 12S rRNA (*12S*), 16S rRNA (*16S*) and tRNAs. All mitochondrial protein-coding genes were free of stop codons and indels, indicating that they were not nuclear insertions. Mitochondrial protein-coding genes were aligned using TranslatorX, while MAFFT (Katoh and Standley 2013) was used to align *12S*, *16S*, and tRNAs. We then concatenated the mitochondrial sequences into 13,393 bp and 14,164 bp alignments, for aphids and *Sulcia*-hosts, respectively.

We tested for substitution saturation using Xia’s method in DAMBE 6 (Xia 2017; Xia and Lemey 2009). Third codon sites in the *Buchnera* (NumOTU = 32, I_SS_ = 0.712, I_SS.C_Asym = 0.601) and aphid data sets were saturated (I_SS_ = 0.4808, I_SS.C_Asym = 0.3932) and thus excluded from our analyses. After we excluded saturated sites, the total lengths of the final data sets were 166,476 bp and 9825 bp (166,416 bp and 8824 bp without outgroups) for the *Buchnera* and mitochondrial alignments, respectively. No evidence of saturation was found in the *Sulcia* data set, but 3rd codon sites, *12S*, and *16S* in the *Sulcia*-host mitochondrial data set were saturated (3rd codon sites: Iss = 0.7218, Iss.cAsym = 0.5570, 12S rRNA: Iss = 0.8932, Iss.cAsym = 0.4332, 16S rRNA: Iss = 0.853, Iss.cAsym = 0.4070) and were excluded from our analyses. After we excluded saturated sites, the total length of the final data set was 8955 bp (8224 bp without outgroups) for the mitochondrial alignment.

### Phylogenetic analysis

RAxML v8.2 (Stamatakis 2014) was used to carry out maximum-likelihood analyses, with 1000 bootstrap replicates to estimate node support. The *Buchnera* data set was partitioned into two subsets: 1st codon sites and 2nd codon sites. The aphid mitochondrial data set was partitioned into four subsets: 1st codon sites, 2nd codon sites, rRNAs, and tRNAs. The *Sulcia* data set was partitioned into three subsets: 1st codon sites, 2nd codon sites, and 3rd codon sites. The *Sulcia*-host mitochondrial data set was partitioned into three subsets: 1st codon sites, 2nd codon sites, and tRNAs. Using jModelTest v2.1.10 (Darriba et al. 2012), we selected the GTR+G substitution model for each data set based on the Bayesian information criterion. We used the Bayesian information criterion in ProtTest v3.4 [13] to select models of the translated amino acids: the CAT+CpREV model for *Buchnera*, the CAT+MtART model for aphid mitochondrial DNA, the CAT+WAG model for *Sulcia*, and the CAT+MtART model for *Sulcia*-host.

We quantified congruence between host and endosymbiont topologies using ParaFit using the R package ape v 5.0 (Paradis and Schliep 2018). We first created matrices of patristic distances calculated from maximum-likelihood phylogenies of host and endosymbiont and a host-endosymbiont association matrix. We then performed a global test with 999 permutations, using the ParafitGlobal value and a *p*-value threshold of 0.05 to determine significance.

### Comparisons of evolutionary rates between hosts and endosymbionts

Root-to-tip distances from the RAxML analyses for each host and endosymbiont pair were calculated and subjected to Pearson correlation analysis using the R packages ape, adephylo (Jombart et al. 2010) and ggpubr (Kassambara 2018). The use of root-to-tip distances removes the confounding effects of time, because all lineages leading to the tips of the tree have experienced the same amount of time since evolving from their common ancestor. However, the sharing of internal branches by groups of taxa renders these data non-independent. Therefore, we compared branch-length differences between hosts and endosymbionts for 15 phylogenetically independent species pairs across aphid and endosymbiont topologies and 14 phylogenetically independent species pairs across *Sulcia* host insects and endosymbiont topologies (see figure S1 and S2). These were calculated using a fixed topology (derived from the endosymbiont analyses described above) for each of the following three data sets: 1) 1st+2nd codon sites of protein-coding genes; 2) translated amino acid sequences; and 3) 1st+2nd codon positions of protein-coding genes plus the inclusion of rRNAs+tRNAs in the case of the aphid mitochondrial data set or the inclusion tRNAs in the case of *Sulcia* host mitochondrial data. Branch lengths were log transformed, and differences between pairs of hosts and pairs of endosymbionts were calculated and compared via Pearson correlation analysis.

To check the data for potential violations of the assumptions of linear regressions, we compared the absolute mean value of log-transformed branch lengths with the log-transformed branch-length differences (Freckleton 2000). We found no significant correlation between these values (*R* = −0.25, *p* = 0.39 for data from aphids; *R* = 0.36, *p* = 0.21 for data from *Buchnera*; *R* = −0.34, *p* = 0.23 for data from *Sulcia* hosts; *R* = 0.15, *p* = 0.61 for data from *Sulcia*), indicating that the data were suitable for use in our analyses. We also performed analyses in which branch-length differences were standardized following previous recommendations (Welch and Waxman 2008), to account for the potentially confounding effects of the different amounts of time that sister-pairs have had to diverge. Three standardizations were carried out, each based on dividing log-transformed branch-length differences by the square root of an estimate of time since divergence for the pair. In the first standardization, time since divergence for host pairs was estimated as the average branch length of the host pair, divided by an assumed rate of 0.001 subs/site/Myr, while for corresponding endosymbionts it was estimated as the average branch length of the endosymbiont pair, divided by the same assumed rate. In the second and third standardizations, times since divergence for both endosymbionts and hosts were based either on average branch lengths of host pairs only or endosymbiont pairs only.

## Results

### Evolutionary rate comparisons between aphids and Buchnera

In all aphid-*Buchnera* analyses, there was strong support for the monophyly of each aphid subfamily and tribe (figures 1, S3 and S4). The topologies inferred from the host and endosymbiont data sets were found to be congruent (*p* = 0.001). In only one case did a disagreement have strong bootstrap support in both trees: *Aphis glycines* is the sister taxon of *Aphis craccivora* in host mitochondrial phylogenies and the sister taxon of *Aphis nasturtii* in endosymbiont phylogenies. This disagreement is likely to be associated with these species being present on the shortest terminal branch lengths in the aphid mitochondrial phylogeny, which could lead to difficulties in resolving their phylogenetic position.

**Fig. 1.**
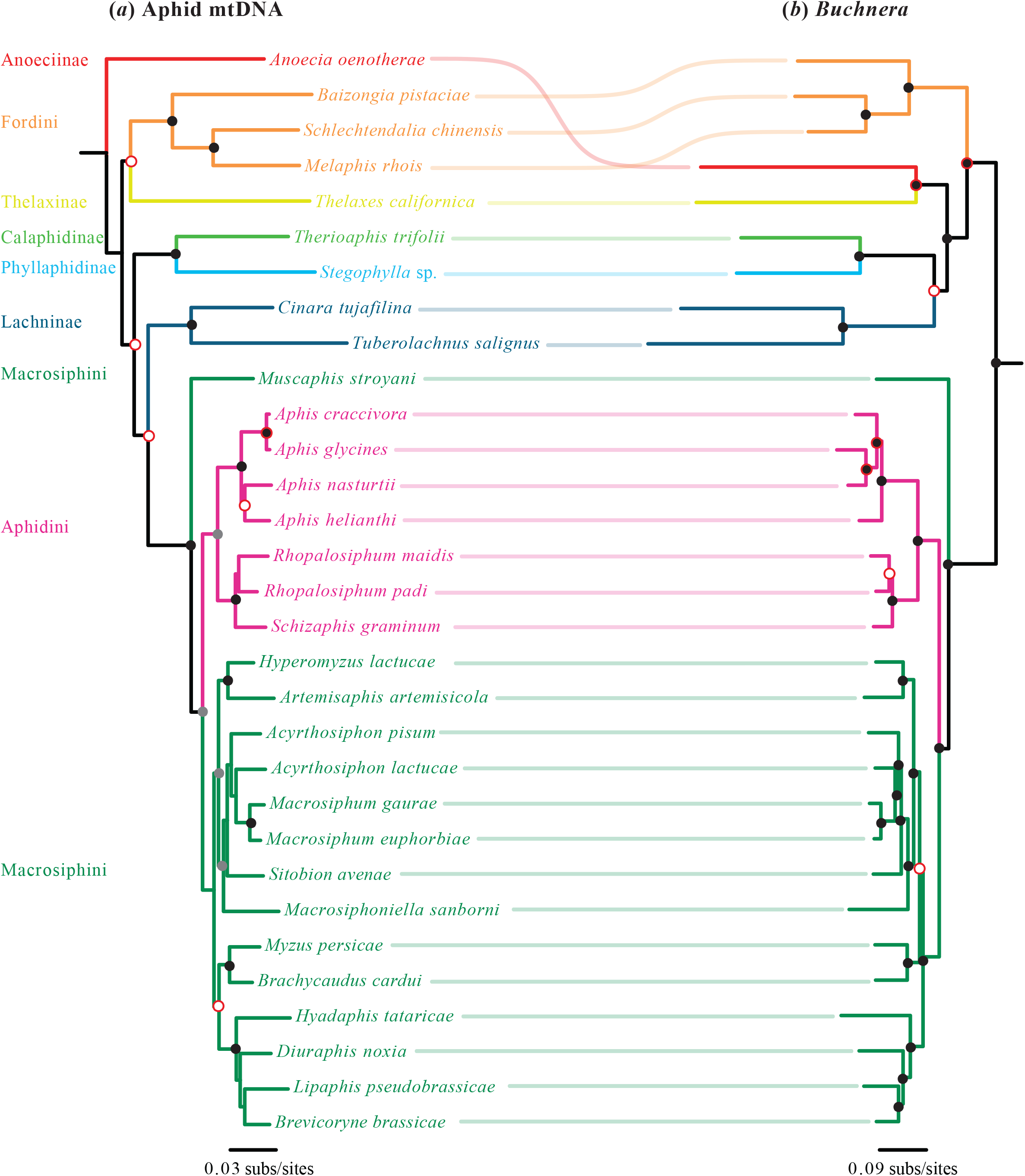
Congruence between (*a*) phylogenetic tree of host aphids inferred using maximum likelihood (RAxML) from whole mitochondrial genomes, and (*b*) phylogenetic tree of *Buchnera* inferred using maximum likelihood from 240 protein-coding genes (3rd codon sites excluded from both data sets). Shaded circles at nodes indicate bootstrap values (black = 100%, grey = 85–99%). Nodes without black or grey circles have bootstrap values <85%. Red outlines on circles indicate disagreement between the phylogenies. Colours of branches indicate different aphid subfamilies or tribes

We found a correlation between root-to-tip distances for data sets including rRNA and tRNA genes in the host data set and protein-coding genes in endosymbiont data set (*R* = 0.88, figure 2a). Similar results were found for data sets including only protein-coding genes from hosts and their endosymbionts (nucleotide alignment: *R*= 0.89, figure S5, amino acid alignment: *R*= 0.91, figure S6). A comparison of branch lengths among phylogenetically independent pairs of host and endosymbiont taxa based on protein-coding genes, including host rRNA and tRNA genes (figure S1) revealed a significant correlation between their rates of evolution (*R* = 0.78, p = 0.0011; figure 2b). A significant correlation was also found when excluding host rRNA and tRNA genes (*R* = 0.77, *p* = 0.013; figure 2c). Equivalent analyses of branch lengths inferred from amino acid data also revealed a significant rate correlation between host and endosymbiont (*R* = 0.83, *p* = 0.0023; figure 2d). Analyses involving standardization of branch-length differences yielded significant rate correlations for all data sets (*R =* 0.72−0.86, *p =* 0.0037−9×10^−5^; see figures S5, S6, and S7)

**Fig. 2.**
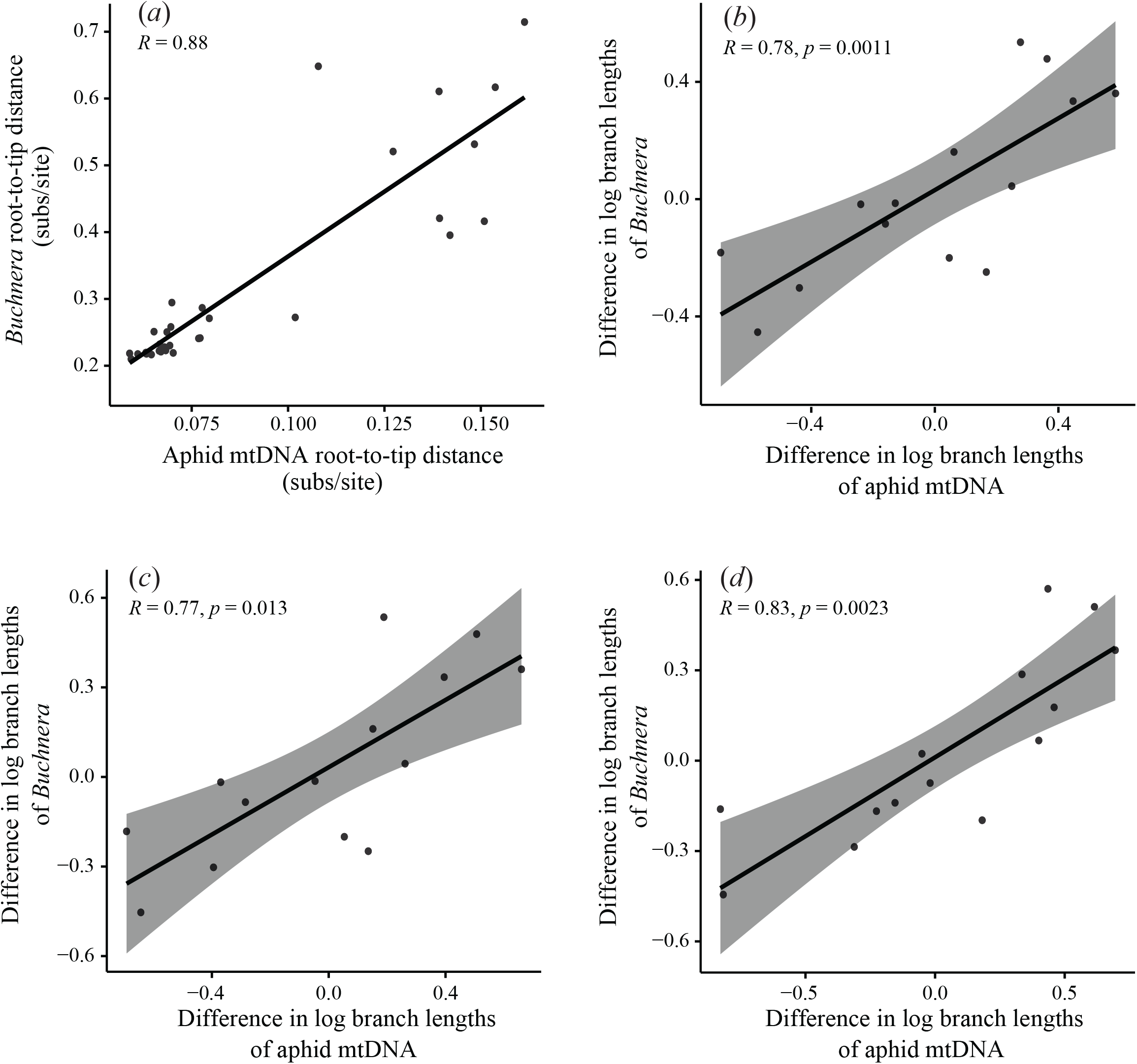
Comparison of evolutionary rates of *Buchnera* symbionts and their aphid hosts. (*a*) Correlation of root-to-tip distances in phylogenies of *Buchnera* and aphids, inferred using maximum-likelihood analysis of whole mitochondrial host genomes and 240 protein-coding *Buchnera* genes (3rd codon sites excluded from both data sets). (*b*) Correlation of log-transformed branch-length differences between phylogenetically independent pairs of host and symbiont taxa, based on whole mitochondrial aphid genomes and 240 protein-coding *Buchnera* genes, (*c*) protein-coding genes only for host mitochondrial data set (excluding 3rd codon sites from both data sets), and (d) amino acid sequences translated from protein-coding genes

### Evolutionary rate comparisons between Sulcia and their hosts

In all analyses of *Sulcia* and their host insects, there was strong support for the monophyly of each insect subfamily, although treehoppers were found to be paraphyletic with respect to leafhoppers (figures 3, S8, and S9). The topologies inferred from all sap-feeding insect mitochondrial DNA and *Sulcia* data sets were congruent (*p* = 0.001). There were no disagreements found to be highly supported in both topologies. We found a correlation between root-to-tip distance for data sets including tRNA genes in the host data set and protein-coding genes in the endosymbiont data set (*R* = 0.92, figure 4). Similar results were found for data sets including only protein-coding genes from hosts and their endosymbionts (nucleotide alignment: *R*= 0.9, figure S10, amino acid alignment: *R*= 0.89, figure S11).

**Fig. 3.**
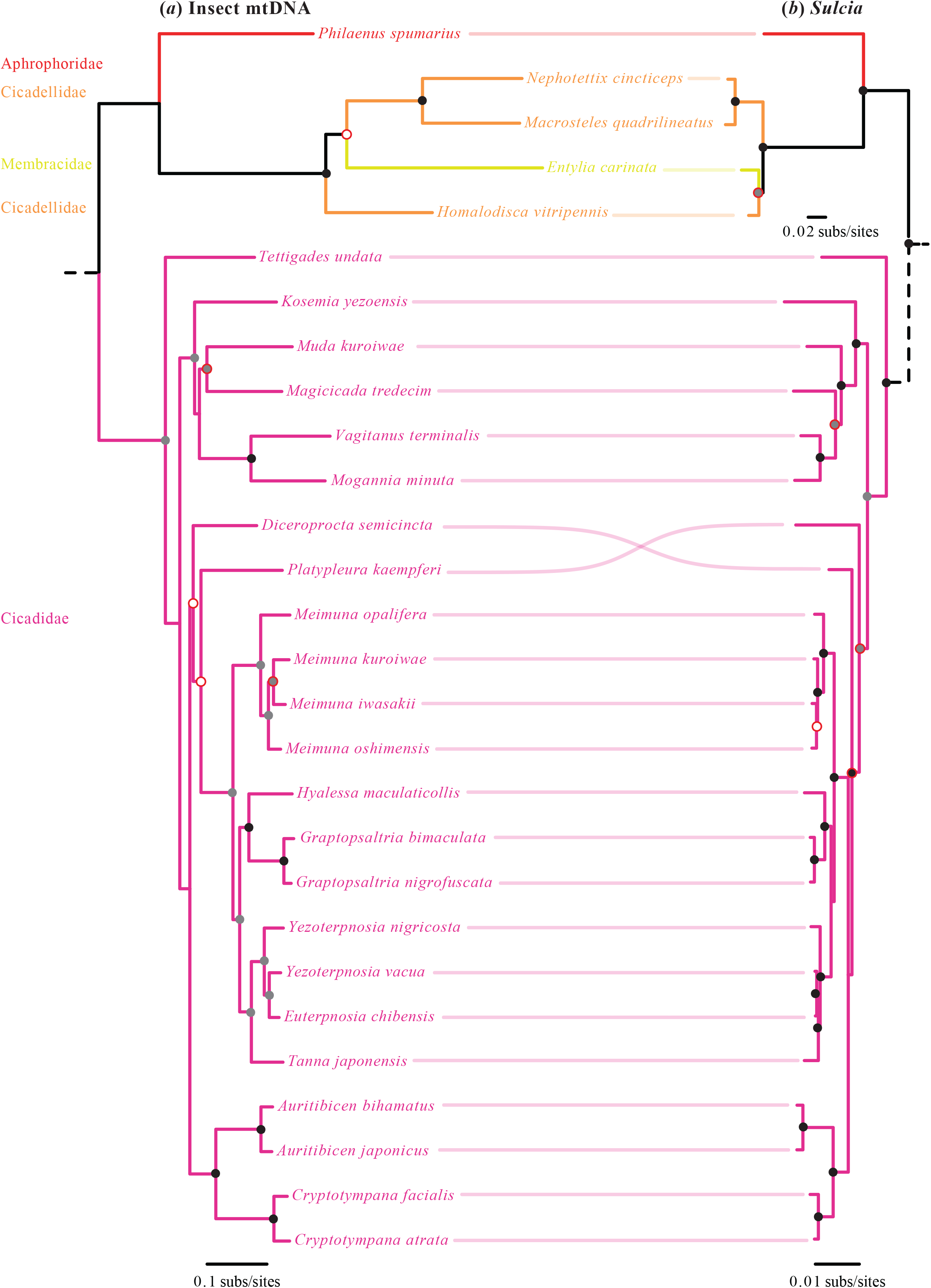
Congruence between (*a*) phylogenetic tree of *Sulcia* hosts inferred using maximum likelihood (RAxML) from mitochondrial protein-coding genes and tRNAs, and (*b*) phylogenetic tree of *Sulcia* inferred using maximum likelihood from 120 protein-coding genes (3rd codon sites excluded from host data set). Shaded circles at nodes indicate bootstrap values (black = 100%, grey = 85−99%). Nodes without black or grey circles have bootstrap values <85%. Red outlines on circles indicate disagreement between the phylogenies. Colours of branches indicate different *Sulcia* host families. Dash dotted branches are not in scale. Solid lines are in scale according to legend underneath them

**Fig. 4.**
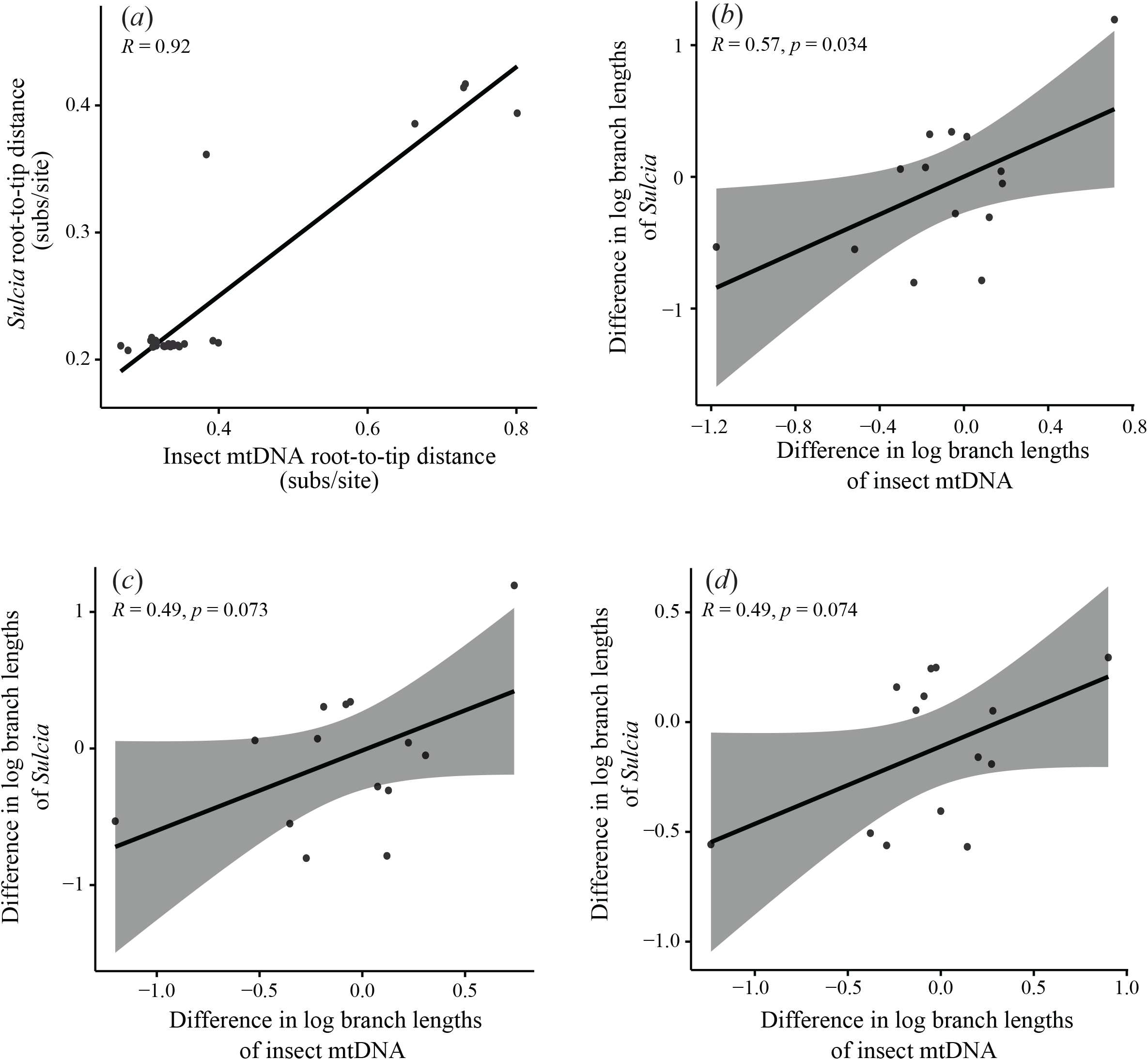
Comparison of evolutionary rates of *Sulcia* symbionts and their hosts. (*a*) Correlation of root-to-tip distances in phylogenies of *Sulcia* and hosts, inferred using maximum-likelihood analysis of host mitochondrial protein-coding genes plus tRNAs and 120 *Sulcia* protein-coding genes, with 3rd codon sites excluded from the host data set. (*b*) Correlation of log-transformed branch-length differences between phylogenetically independent pairs of host and symbiont taxa, based on mitochondrial protein-coding genes plus tRNAs for the host data set, (*c*) exclusion of 3rd codon sites and tRNAs from the host mitochondrial data set, and (d) amino acid sequences translated from protein-coding genes

A comparison of branch lengths among phylogenetically independent pairs of host and endosymbiont taxa based on protein-coding genes, including host tRNA genes (figure S2) revealed a significant correlation between their rates of evolution (*R* = 0.62, p = 0.017; figure 4b). However, a non-significant correlation was found when excluding host tRNAs (*R* = 0.49, *p* = 0.073; figure 4c). Equivalent analyses of branch lengths inferred from amino acid data also revealed a weak non-significant rate correlation between host and endosymbiont (*R* = 0.49, *p* = 0.074; figure 4d). Analyses involving standardization of branch-length differences yielded non-significant, weak rate correlations for all data sets (*R =* 0.054–0.35, *p =* 0.17–0.85; see figures S10, S11 and S12).

## Discussion

We have detected a correlation in molecular evolutionary rates between *Buchnera* and host mitochondrial genomes, using two different methods of analysis. On the other hand, a correlation in molecular evolutionary rates was detected between *Sulcia* and host mitochondrial genomes using the same methods, although it was not statistically significant. Our previous work found similar correlations in the evolutionary rates of cockroach mitochondrial genomes and the genomes of their endosymbionts *Blattabacterium* (Arab et al. 2020). Other studies have found a correlation in evolutionary rates between nuclear and mitochondrial genes in various animals (Lourenço et al. 2013; Martin 1999; Sheldon et al. 2000; Yan et al. 2019), between mitochondrial and plastid genes in angiosperms (Sloan et al. 2012), and between nuclear, mitochondrial and plastid genes in algae (Hua et al. 2012; Smith and Lee 2010).

A relationship between host and endosymbiont substitution rates could be explained by similar forces acting on their underlying mutation rates. One way in which this could occur is if endosymbiont DNA replication depends on the host’s DNA replication and repair machinery (Moran and Bennett 2014). Some *Buchnera* genomes show a reduction in the number of repair and replication genes, which suggests that the endosymbiont might be dependent on host DNA repair and replication machinery, potentially leading to similarities in their mutation rates (Moran and Bennett 2014; Silva et al. 2001). Correlations in evolutionary rates could also be the result of interactions between host mitochondrial and endosymbiont proteins, as has been found between insect mitochondrial-encoded and nuclear-encoded OXPHOS genes associated with mitochondrial function (Yan et al. 2019). There is evidence of close physical and metabolic association between *Buchnera* and mitochondria. Electron microscopy studies indicate a dense number of mitochondria in aphid bacteriocytes (Griffiths and Beck 1973; Hinde 1971). A transcriptomic study of pea aphid (*Acyrthosiphon pisum*) found an upregulation of several genes for mitochondria-related transporters in bacteriocytes, suggesting cooperative metabolic interactions between *Buchnera* and mitochondria (Nakabachi et al. 2005). A close physical and metabolic association between *Buchnera* and mitochondria indicates that changes affecting mitochondrial genome evolution could impact the genome of the endosymbiont, explaining the correlation in molecular rate between the two. The extent of interactions between host and *Buchnera* proteins, however, remains to be investigated.

A significant correlation was not detected when comparing evolutionary rates between *Sulcia* and host mitochondrial genomes. This result contrasted with those of analyses using *Blattabacterium*, *Buchnera* genomes and mitochondrial genomes of their hosts. One explanation for this result could be the number of samples included in the data set from different host families. Taxa that were included in these analyses were limited to members of only four families of insects, with 22 out of 28 species belonging to Cicadidae. Additionally, those 22 samples came from one geographical location, Japan. Future studies on evolutionary rate correlations between *Sulcia* and host mitochondria should include more representatives of Auchenorrhyncha that contain *Sulcia* to test the hypothesis that their evolutionary rates are not correlated.

To further explain why a significant correlation in evolutionary rates was not detected between *Sulcia* and their hosts, it is important to consider the biology and evolutionary history of *Sulcia*. It is believed that the ancestor of Auchenorrhyncha acquired Sulcia 260 Mya, making it the oldest known insect endosymbiont. Genomes of *Sulcia* are highly reduced and are among the smallest of all known insect endosymbionts. These tiny genomes lack genes responsible for cellular information processing mechanisms, which are essential for DNA repair, replication, and translation. *Sulcia* is always found in hosts with at least one other co-primary endosymbiont (Bennett and Moran 2013). Co-primary endosymbionts have been lost and replaced multiple times in Auchenorrhyncha. For example, *Candidatus* Nasuia deltocephalinicola (herafter *Nasuia*) is believed to be an ancient endosymbiont that probably infected the ancestor of Auchenorrhyncha (Bennett and Moran 2013), while *Candidatus* Baumannia cicadellinicola (hereafter *Baumannia*) is believed to have replaced *Nasuia* more recently in the ancestor of sharpshooters (McCutcheon and Moran 2007).

A recent study found a lack of overexpression of several host repair and replication genes that might replace these functions in the endosymbiont in the glassy-winged sharpshooter (*Homalodisca vitripennis*) (Mao and Bennett 2020). An alternative source that may assist *Sulcia* with these functions is their co-primary endosymbionts. Genomes of co-primary endosymbionts that are found in *Sulcia* hosts vary in size. *Nausia* has the smallest known genome of any insect endosymbiont (Bennett and Moran 2013), while *Baumannia* has a relatively large genome (McCutcheon and Moran 2007). A transcriptomic study found an overexpression of host genes in the bacteriocytes of the aster leafhopper (*Macrosteles quadrilineatus*) that are believed to provide support to *Sulcia* and *Nasuia* by performing functions such as replication, transcription, and translation (Mao et al. 2018). However, while investigating glassy-winged sharpshooter bacteriocytes containing *Sulcia*, Mao and Bennett (2020) found that several genes that are responsible for DNA replication and repair were missing. An analysis of the genome of *Baumannia* found large numbers of repair and replication genes in comparison with other known insect endosymbionts (Wu et al. 2006). In this case, *Sulcia* might depend on *Baumannia* instead of the host’s for DNA replication and repair in the glassy-winged sharpshooter, but it is unknown whether this occurs between insect co-primary endosymbionts.

The non-significant correlation between *Sulcia* and host mitochondrial genomes evolutionary rates in this study could be due to representative hosts having co-primary endosymbionts with relatively large genomes within *Sulcia* hosts. Sixteen out of 28 species examined in this study harbour a co-primary endosymbiont with a large genome; in addition to *Sulcia*, 14 cicada species contain a yeast-like endosymbiont (25.1 Mb) (Matsuura et al. 2018), while *H. vitripennis* harbours *Baumannia* (686 kb) (Wu et al. 2006) and *Philaenus spumarius* harbours an endosymbiont closely related to *Sodalis glossinidius* (4.1 Mb) (Koga and Moran 2014). This indicates an overrepresentation of *Sulcia* hosts with recent co-primary endosymbionts with relatively large genomes that presumably retain function. If co-primary endosymbionts with complete DNA repair and replication machinery were found to provide those essential replication and repair services to *Sulcia*, then this lack of dependence on host DNA repair and replication machinery could influence the evolutionary rate correlation between host and endosymbiont genomes. We now know that hosts of *Sulcia* have evolved tailored mechanisms to support each co-primary endosymbiont. To that end, future studies should investigate correlations of evolutionary rates between genomes of *Sulcia*, co-primary endosymbionts with large genomes, and their hosts.

*Buchnera* and *Sulcia* are vertically transmitted, obligate intercellular mutualistic endosymbionts, whose phylogeny is expected to mirror that of their hosts. This is expected for phylogenies inferred from mitochondrial DNA, since mitochondria are linked with the endosymbionts through vertical transfer to offspring through the egg cytoplasm. As observed in previous studies (Clark et al. 2000; Nováková et al. 2013; Urban and Cryan 2012), we found a high level of agreement between the topologies inferred from aphid mitochondrial and *Buchnera* genome data sets and the same was found for host mitochondrial and *Sulcia* genome data sets. In the case of aphids and *Buchnera*, however, we observed disagreements between some well-supported relationships. The variability in rates observed between some lineages, coupled with the highly increased rate of mitochondrial DNA compared with *Buchnera* DNA and very short terminal branches of some taxa on host mitochondrial phylogenies, could be responsible for these disagreements. Regarding insect host mitochondrial and *Sulcia* genome data sets, some disagreements were observed between relationships with low support in either topology. This could be explained by the highly increased rate of mitochondrial DNA compared with *Sulcia* DNA.

In conclusion, our results for the aphid-*Buchnera* data set highlight the significant effects that long-term symbiosis can have on the biology of each symbiotic partner. The rate of evolution is a fundamental characteristic of any species; our study indicates that it can become closely linked between organisms as a result of symbiosis. Detecting correlations in evolutionary rates between hosts and endosymbionts can highlight forces impacting the evolution of both organisms. We provide evidence that these correlations extend to multiple groups of insects and their endosymbionts. To further our understanding of the extent of these correlations, future work is required to determine whether the correlation that we have found here also applies to the nuclear genome of the host. Additionally, analyses performed in this work should be applied to different groups of hosts and their endosymbionts to assess how widespread these correlations are between the genomes of the two. On the other hand, our results for the *Sulcia* endosymbiont and the mitochondrial genomes of its hosts did not show a significant correlation in evolutionary rates. Further studies should examine whether this result is due to limitations in the data set used, or it is a result of acquisition co-primary endosymbionts by the insect hosts. This can highlight the impact of acquiring new co-primary endosymbionts on the evolution of the genomes of both host and ancient endosymbionts.

## Declarations

### Ethics approval

Not Applicable

### Consent to Participate

Not applicable.

### Consent for publication

All authors consent for publishing this manuscript.

### Availability of data and material

Datasets will be uploaded to DRYAD repository after submitting the manuscript.

### Code availability

Not applicable

### Author’s contribution

All authors contributed to the study conception and design. Material preparation, data collection and analysis were performed by Daej A Arab. The first draft of the manuscript was written by Daej A Arab and all authors commented on previous versions of the manuscript. All authors read and approved the final manuscript.

## References

Abascal F, Zardoya R, Telford MJ (2010) TranslatorX: multiple alignment of nucleotide sequences guided by amino acid translations. Nucleic Acids Res 38:W7–W13

Altenhoff AM, Levy J, Zarowiecki M, Tomiczek B, Vesztrocy AW, Dalquen DA, Müller S, Telford MJ, Glover NM, Dylus D (2019) OMA standalone: orthology inference among public and custom genomes and transcriptomes. Genome Res 29:1152–1163

Arab DA, Bourguignon T, Wang Z, Ho SY, Lo N (2020) Evolutionary rates are correlated between cockroach symbionts and mitochondrial genomes. Biol Lett 16:20190702

Baumann P (2005) Biology of bacteriocyte-associated endosymbionts of plant sap-sucking insects. Annu Rev Microbiol 59:155–189

Baumann P, Baumann L, Lai C-Y, Rouhbakhsh D, Moran NA, Clark MA (1995) Genetics, physiology, and evolutionary relationships of the genus Buchnera: intracellular symbionts of aphids. Annu Rev Microbiol 49:55–94

Bennett GM, Moran NA (2013) Small, smaller, smallest: the origins and evolution of ancient dual symbioses in a phloem-feeding insect. Genome Biol Evol 5:1675–1688

Bernt M, Donath A, Jühling F, Externbrink F, Florentz C, Fritzsch G, Pütz J, Middendorf M, Stadler PF (2013) MITOS: improved de novo metazoan mitochondrial genome annotation. Mol Phylogenet Evol 69:313–319

Braendle C, Miura T, Bickel R, Shingleton AW, Kambhampati S, Stern DL (2003) Developmental origin and evolution of bacteriocytes in the aphid–Buchnera symbiosis. PLoS Biol 1:e21

Bromham L (2009) Why do species vary in their rate of molecular evolution? Biol Lett 5:401–404

Bromham L, Penny D (2003) The modern molecular clock. Nat Rev Genet 4:216–224

Chong RA, Moran NA (2018) Evolutionary loss and replacement of Buchnera, the obligate endosymbiont of aphids. ISME J 12:898–908

Clark MA, Moran NA, Baumann P, Wernegreen JJ (2000) Cospeciation between bacterial endosymbionts (Buchnera) and a recent radiation of aphids (Uroleucon) and pitfalls of testing for phylogenetic congruence. Evolution 54:517–525

Darriba D, Taboada GL, Doallo R, Posada D (2012) jModelTest 2: more models, new heuristics and parallel computing. Nat Methods 9:772

Douglas A (1998) Nutritional interactions in insect-microbial symbioses: aphids and their symbiotic bacteria Buchnera. Annu Rev Entomol 43:17–37

Douglas AE (2010) The Symbiotic Habit. Princeton University Press, Princeton, NJ

Freckleton RP (2000) Phylogenetic tests of ecological and evolutionary hypotheses: checking for phylogenetic independence. Funct Ecol 14:129–134

Fukatsu T, Ishikawa H (1996) Phylogenetic position of yeast-like symbiont of Hamiltonaphis styraci (Homoptera, Aphididae) based on 18S rDNA sequence. Insect Biochem Mol Biol 26:383–388

Griffiths GW, Beck SD (1973) Intracellular symbiotes of the pea aphid, Acyrthosiphon pisum. J Insect Physiol 19:75–84

Hansen AK, Moran NA (2011) Aphid genome expression reveals host–symbiont cooperation in the production of amino acids. Proc Natl Acad Sci 108:2849–2854

Hinde R (1971) The control of the mycetome symbiotes of the aphids Brevicoryne brassicae, Myzus persicae, and Macrosiphum rosae. J Insect Physiol 17:1791–1800

Ho SYW, Lo N (2013) The insect molecular clock. Aust J Entomol 52:101–105

Hua J, Smith DR, Borza T, Lee RW (2012) Similar relative mutation rates in the three genetic compartments of Mesostigma and Chlamydomonas. Protist 163:105–115

Jombart T, Balloux F, Dray S (2010) Adephylo: new tools for investigating the phylogenetic signal in biological traits. Bioinformatics 26:1907–1909

Kassambara A (2018) ggpubr:“ggplot2” based publication ready plots. R package version 0.2.

Katoh K, Standley DM (2013) MAFFT multiple sequence alignment software version 7: improvements in performance and usability. Mol Biol Evol 30:772–780

Koga R, Meng X-Y, Tsuchida T, Fukatsu T (2012) Cellular mechanism for selective vertical transmission of an obligate insect symbiont at the bacteriocyte–embryo interface. Proc Natl Acad Sci 109:E1230–E1237

Koga R, Moran NA (2014) Swapping symbionts in spittlebugs: evolutionary replacement of a reduced genome symbiont. ISME J 8:1237–1246

Lourenço JM, Glémin S, Chiari Y, Galtier N (2013) The determinants of the molecular substitution process in turtles. J Evol Biol 26:38–50

Mao M, Bennett GM (2020) Symbiont replacements reset the co-evolutionary relationship between insects and their heritable bacteria. ISME J:1–12

Mao M, Yang X, Bennett GM (2018) Evolution of host support for two ancient bacterial symbionts with differentially degraded genomes in a leafhopper host. Proc Natl Acad Sci 115:E11691–E11700

Martin AP (1999) Substitution rates of organelle and nuclear genes in sharks: implicating metabolic rate (again). Mol Biol Evol 16:996–1002

Matsuura Y, Moriyama M, Łukasik P, Vanderpool D, Tanahashi M, Meng X-Y, McCutcheon JP, Fukatsu T (2018) Recurrent symbiont recruitment from fungal parasites in cicadas. Proc Natl Acad Sci 115:E5970–E5979

McCutcheon JP, Moran NA (2007) Parallel genomic evolution and metabolic interdependence in an ancient symbiosis. Proc Natl Acad Sci 104:19392–19397

McCutcheon JP, Moran NA (2010) Functional convergence in reduced genomes of bacterial symbionts spanning 200 My of evolution. Genome Biol Evol 2:708–718

Moran NA, Bennett GM (2014) The tiniest tiny genomes. Annu Rev Microbiol 68:195–215

Moran NA, McCutcheon JP, Nakabachi A (2008) Genomics and evolution of heritable bacterial symbionts. Annu Rev Genet 42:165–190

Moran NA, Munson MA, Baumann P, Ishikawa H (1993) A molecular clock in endosymbiotic bacteria is calibrated using the insect hosts. Proc Royal Soc B 253:167–171

Moran NA, Tran P, Gerardo NM (2005) Symbiosis and insect diversification: an ancient symbiont of sap-feeding insects from the bacterial phylum Bacteroidetes. Appl Environ Microbiol 71:8802–8810

Nakabachi A, Shigenobu S, Sakazume N, Shiraki T, Hayashizaki Y, Carninci P, Ishikawa H, Kudo T, Fukatsu T (2005) Transcriptome analysis of the aphid bacteriocyte, the symbiotic host cell that harbors an endocellular mutualistic bacterium, Buchnera. Proc Natl Acad Sci 102:5477–5482

Nováková E, Hypša V, Klein J, Foottit RG, von Dohlen CD, Moran NA (2013) Reconstructing the phylogeny of aphids (Hemiptera: Aphididae) using DNA of the obligate symbiont Buchnera aphidicola. Mol Phylogenet Evol 68:42–54

Paradis E, Schliep K (2018) ape 5.0: an environment for modern phylogenetics and evolutionary analyses in R. Bioinformatics 35:526–528

Sheldon FH, Jones CE, McCracken KG (2000) Relative patterns and rates of evolution in heron nuclear and mitochondrial DNA. Mol Biol Evol 17:437–450

Silva FJ, Latorre A, Moya A (2001) Genome size reduction through multiple events of gene disintegration in Buchnera APS. Trends Genet 17:615–618

Sloan DB, Alverson AJ, Wu M, Palmer JD, Taylor DR (2012) Recent acceleration of plastid sequence and structural evolution coincides with extreme mitochondrial divergence in the angiosperm genus Silene. Genome Biol Evol 4:294–306

Smith DR, Lee RW (2010) Low nucleotide diversity for the expanded organelle and nuclear genomes of Volvox carteri supports the mutational-hazard hypothesis. Mol Biol Evol 27:2244–2256

Stamatakis A (2014) RAxML version 8: a tool for phylogenetic analysis and post-analysis of large phylogenies. Bioinformatics 30:1312–1313

Urban JM, Cryan JR (2012) Two ancient bacterial endosymbionts have coevolved with the planthoppers (Insecta: Hemiptera: Fulgoroidea). BMC Evol Biol 12:87

Welch JJ, Waxman D (2008) Calculating independent contrasts for the comparative study of substitution rates. J Theor Biol 251:667–678

Wu D, Daugherty SC, Van Aken SE, Pai GH, Watkins KL, Khouri H, Tallon LJ, Zaborsky JM, Dunbar HE, Tran PL (2006) Metabolic complementarity and genomics of the dual bacterial symbiosis of sharpshooters. PLoS Biol 4:e188

Xia X (2017) DAMBE6: new tools for microbial genomics, phylogenetics, and molecular evolution. J Hered 108:431–437

Xia X, Lemey P (2009) Assessing substitution saturation with DAMBE. In: Lemey P, Salemi M, Vandamme A-M (eds) The phylogenetic handbook: a practical approach to DNA and protein phylogeny. Cambridge University Press, Cambridge, pp. 615–630

Yan Z, Ye G, Werren JH (2019) Evolutionary rate correlation between mitochondrial-encoded and mitochondria-associated nuclear-encoded proteins in insects. Mol Biol Evol 36:1022–1036

Zerbino DR (2010) Using the velvet de novo assembler for short-read sequencing technologies. Curr Protoc Bioinformatics 31:11.5.1–11.5.12

