## Supplementary Figures for "Evolutionary rates are correlated between *Buchnera* endosymbionts and mitochondrial genomes of their aphid hosts"

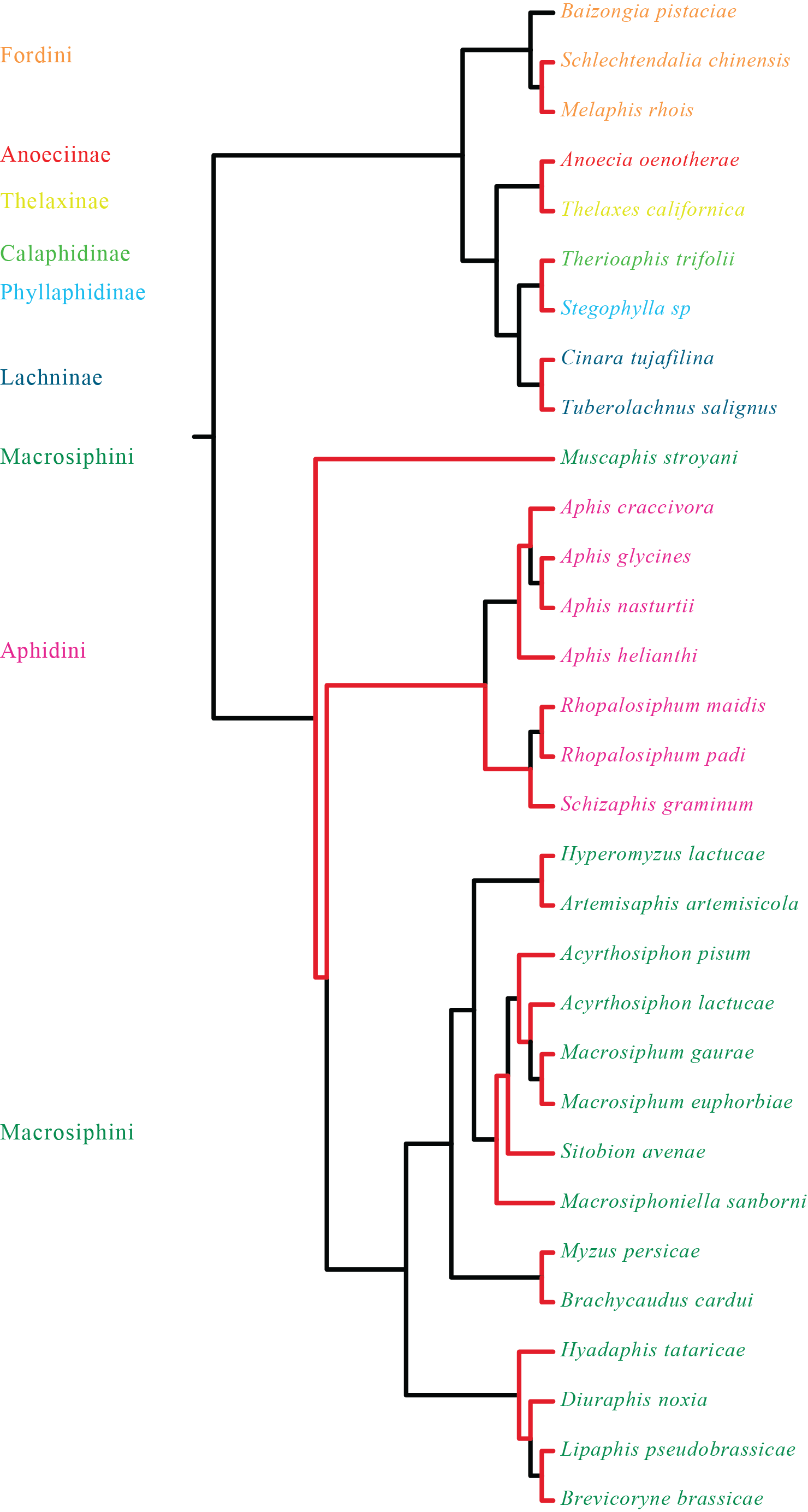

**Fig. S1** A fixed tree topology (obtained from the *Buchnera* tree shown in Fig. 1) used in all analyses of root-to-tip distances and branch-length comparisons. Fifteen phylogenetically independent pairs of lineages used to test for correlations of evolutionary rates are shown in red. Species names are coloured according to the family or tribe to which they belong, as shown on the left of the figure

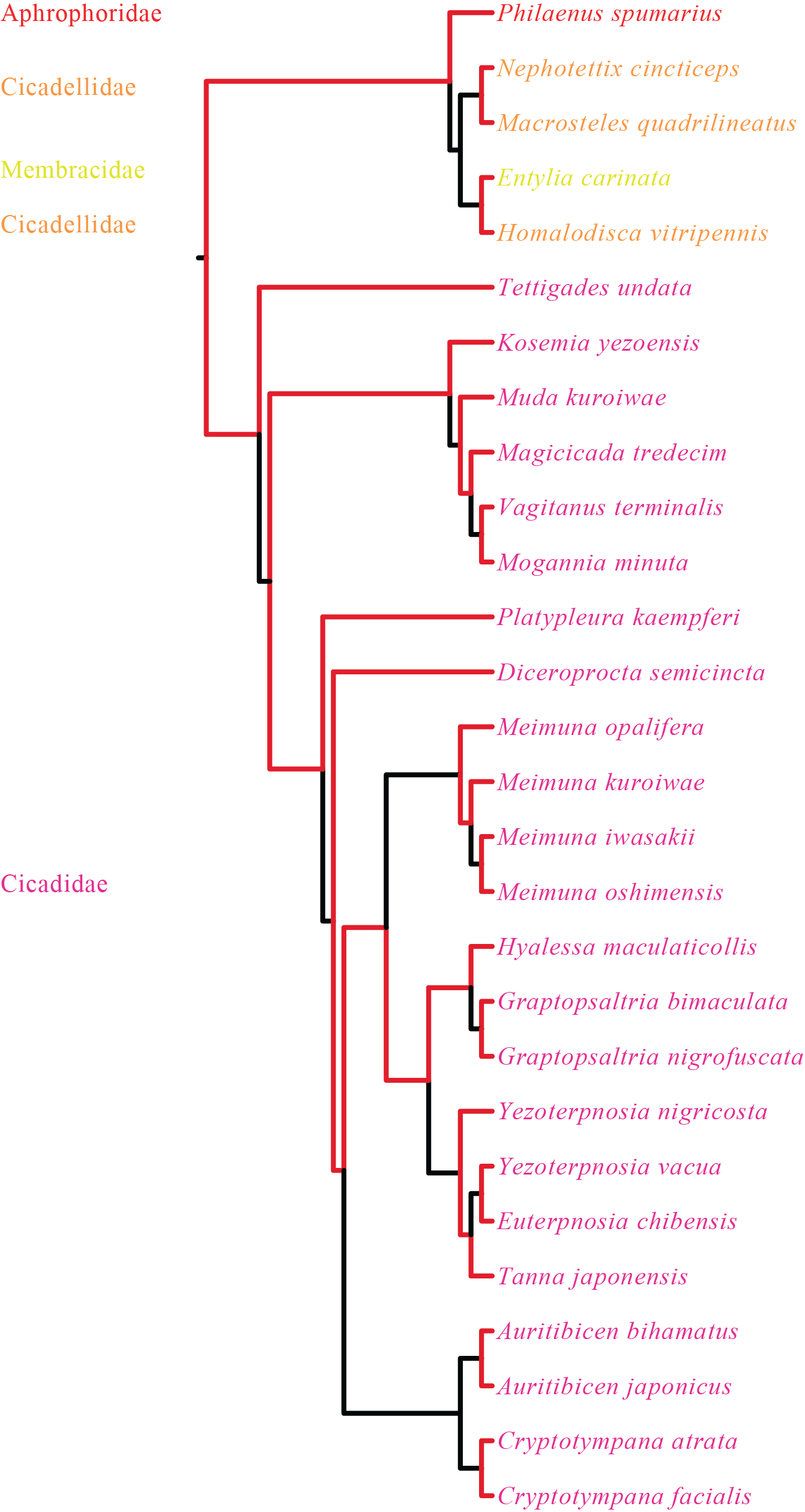

**Fig. S2** A fixed tree topology (obtained from the *Sulcia* tree shown in Fig. 3) was used in all analyses of root-to-tip distances and branch-length comparisons. Fourteen phylogenetically independent pairs of lineages used to test for correlations of evolutionary rates are shown in red. Species names are coloured according to the family to which they belong, as shown on the left of the figure

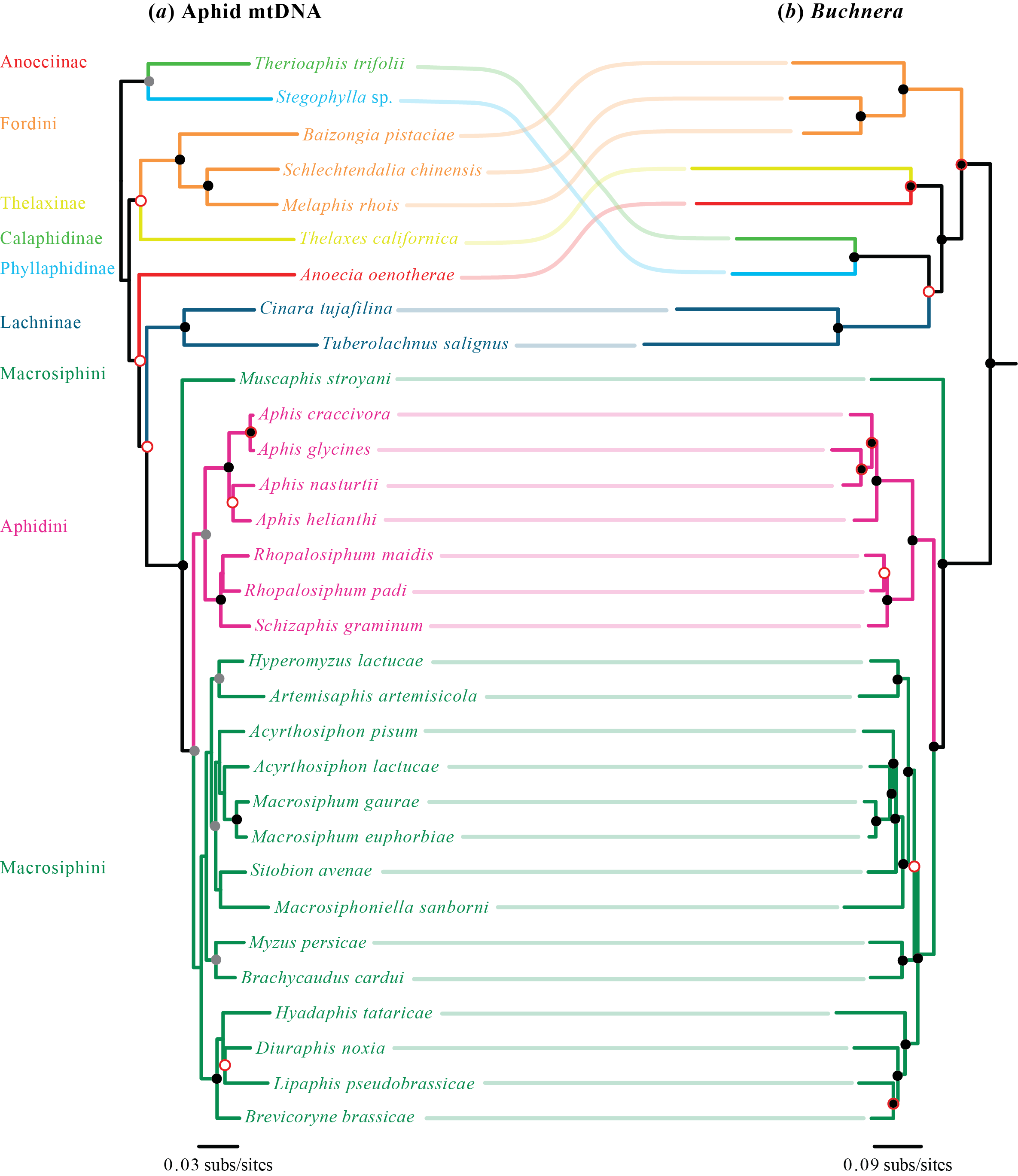

**Fig. S3** Congruence between (*a*) phylogenetic tree of host aphids inferred using maximum likelihood (RAxML) from mitochondrial protein-coding genes, and (*b*) phylogenetic tree of *Buchnera* inferred using maximum likelihood from 240 protein-coding genes (3rd codon sites excluded from both data sets). Shaded circles at nodes indicate bootstrap values (black = 100%, grey = 85–99%). Nodes without black or grey circles have bootstrap values <85%. Red outlines on circles indicate disagreement between the phylogenies. Colours of branches indicate different aphid families

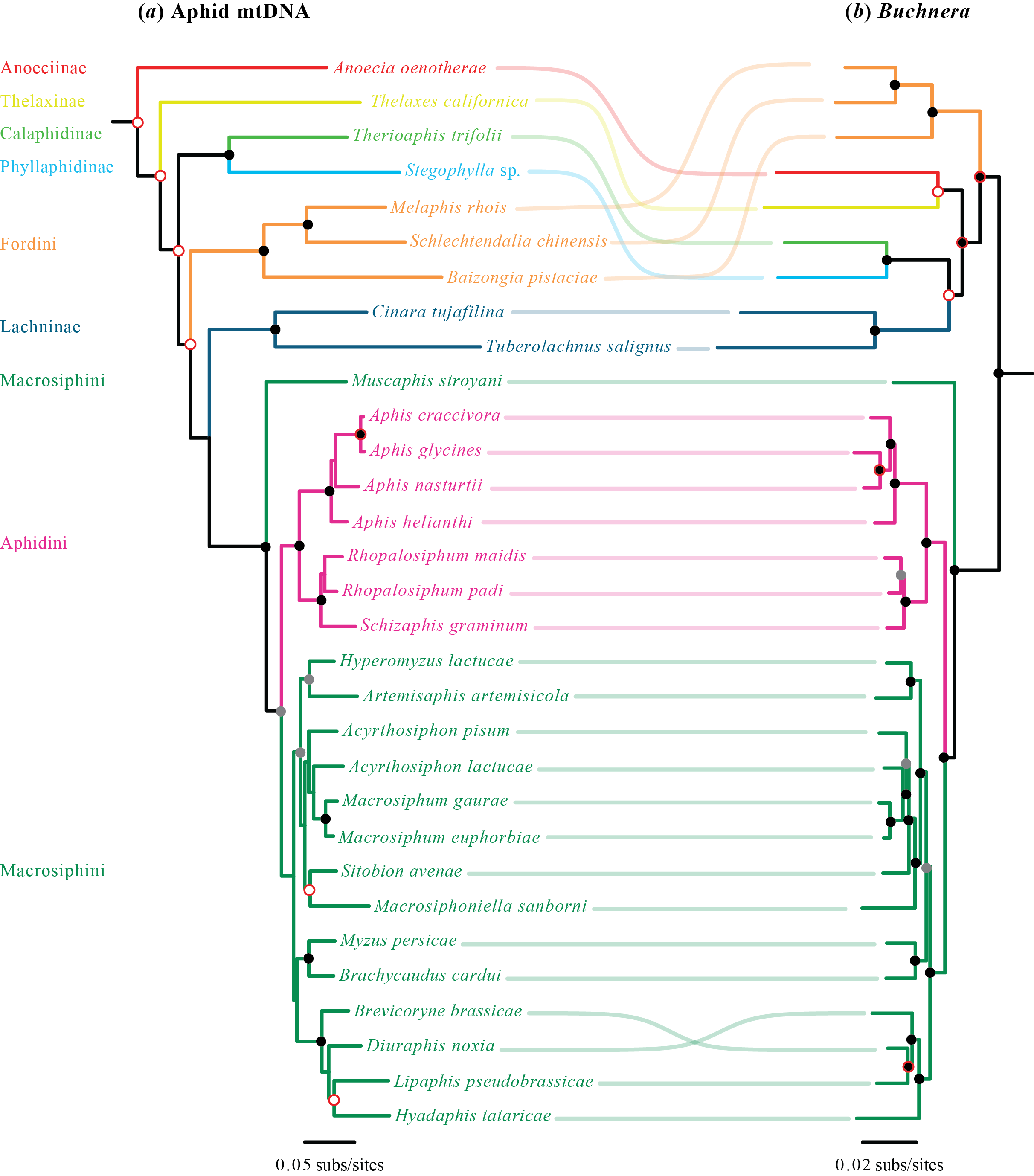

**Supplementary Fig. S4** Congruence between phylogenetic trees inferred using maximum likelihood (RAxML) from translated amino acid data sets of (*a*) mitochondrial protein-coding genes from host aphids, and (*b*) 240 protein-coding genes from *Buchnera*. Shaded circles at nodes indicate bootstrap values (black = 100%, grey = 85–99%). Nodes without black or grey circles have bootstrap values <85%. Red outlines on circles indicate disagreement between the phylogenies. Colours of branches indicate different aphid families

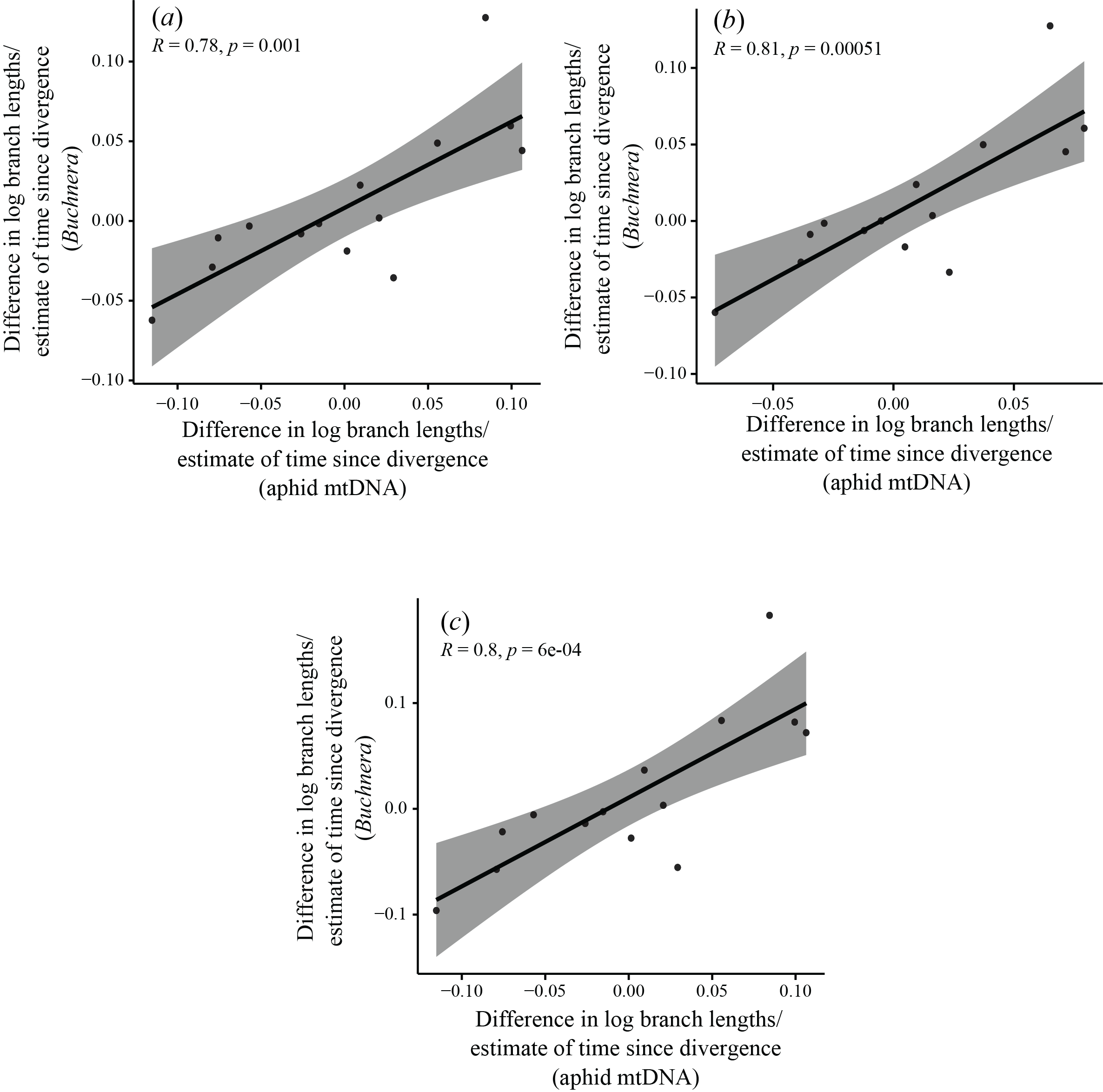

**Fig. S5** Standardized tests for correlation of evolutionary rates between 15 independent pairs of *Buchnera* and host aphid mitochondria, based on 240 symbiont genes and whole host mitochondrial genomes, with 3rd codon sites excluded from both data sets. Three standardizations were carried out, each based on dividing log-transformed branch-length differences by the square root of an estimate of time since divergence for the pair. In the first standardization (*a*), time since divergence for host pairs was estimated as the average branch length of the host pair, divided by an assumed rate of 0.001 subs/site/Myr, while for corresponding symbionts it was estimated as the average branch length of the symbiont pair, divided by the same assumed rate. In the second (*b*) and third (*c*) standardizations, times since divergence for both symbionts and hosts were based either on average branch lengths of host pairs only or symbiont pairs only

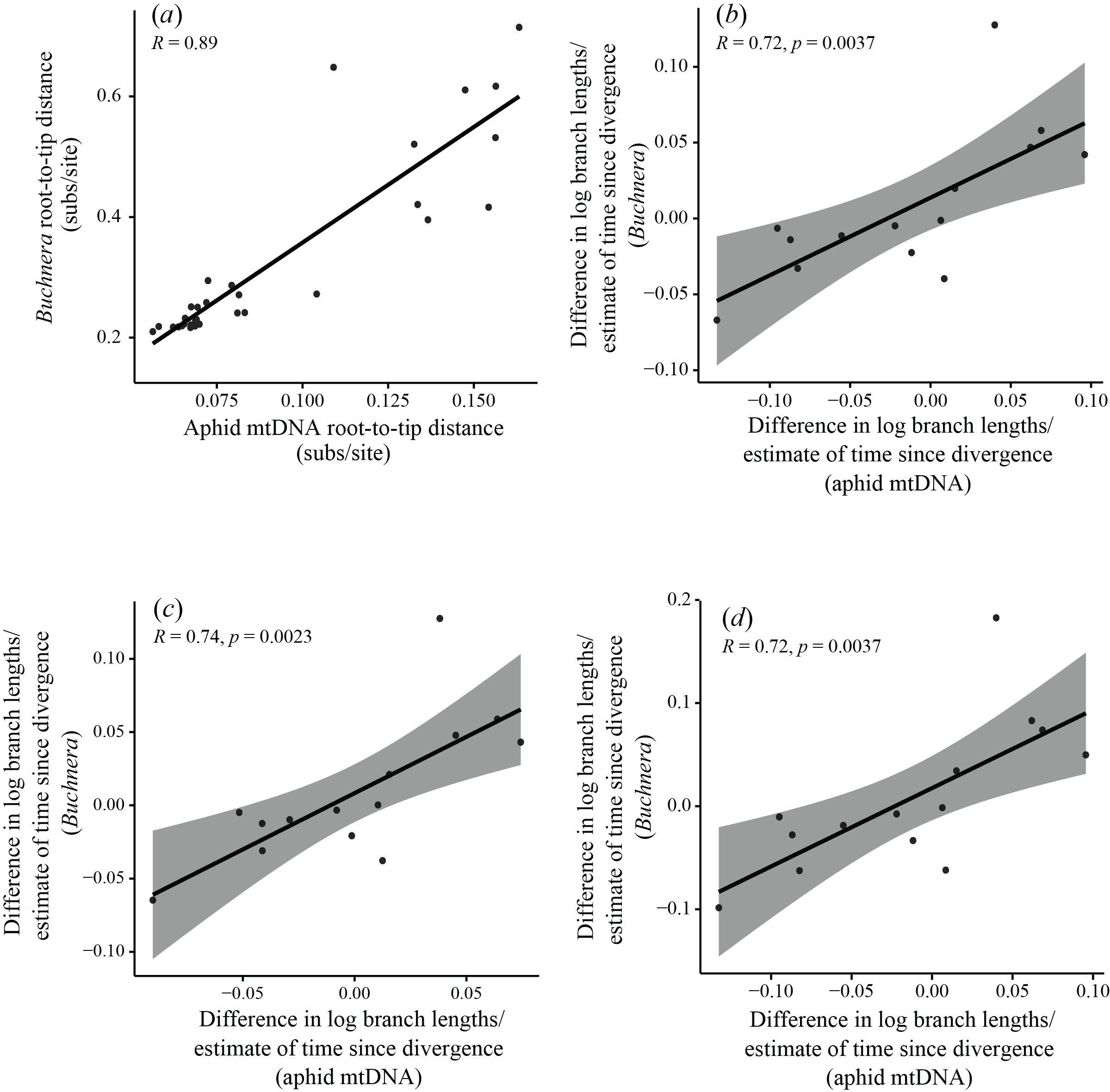

**Supplementary Fig. S6** Comparison of evolutionary rates of *Buchnera* symbionts and their aphid hosts. (*a*) Correlation of root-to-tip distances in phylogenies of *Buchnera* and aphids, inferred using maximum-likelihood analysis of protein-coding genes from each data set, with 3rd codon sites excluded. (*b–d*) Standardized tests for correlation of molecular evolutionary rates between 15 independent pairs of *Buchnera* and host aphid mitochondria. Three standardizations were carried out, each based on dividing log-transformed branch-length differences by the square root of an estimate of time since divergence for the pair. In the first (*b*), time since divergence for host pairs was estimated as the average branch length of the host pair, divided by an assumed rate of 0.001 subs/site/million years, while for corresponding symbionts it was estimated as the average branch length of the symbiont pair, divided by the same assumed rate. In the second (*c*) and third (*d*) standardizations, times since divergence for both symbionts and hosts were based either on average branch lengths of host pairs only or symbiont pairs only

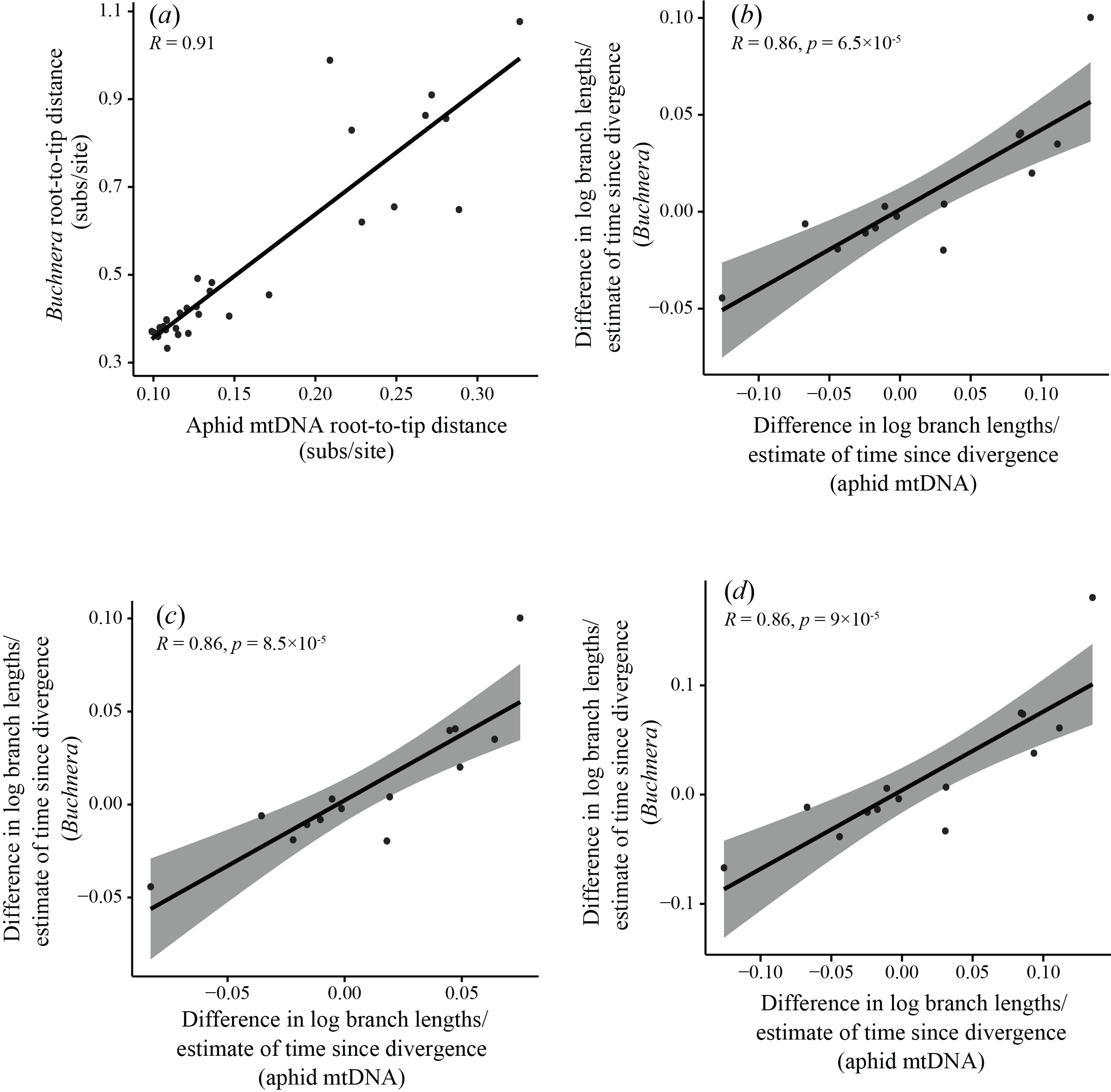

**Supplementary Fig. S7** Comparison of evolutionary rates of *Buchnera* symbionts and their aphid hosts. (*a*) Correlation of root-to-tip distances in phylogenies of *Buchnera* and aphids, inferred using maximum-likelihood analysis of amino acid sequences translated from protein-coding genes. (*b–d*) Standardized tests for correlation of molecular evolutionary rates between 15 independent pairs of *Buchnera* and host aphid mitochondria. Three standardizations were carried out, each based on dividing log-transformed branch-length differences by the square root of an estimate of time since divergence for the pair. In the first standardization (*b*), time since divergence for host pairs was estimated as the average branch length of the host pair, divided by an assumed rate of 0.001 subs/site/million years, while for corresponding symbionts it was estimated as the average branch length of the symbiont pair, divided by the same assumed rate. In the second (*c*) and third (*d*) standardizations, times since divergence for both symbionts and hosts were based either on average branch lengths of host pairs only or symbiont pairs only

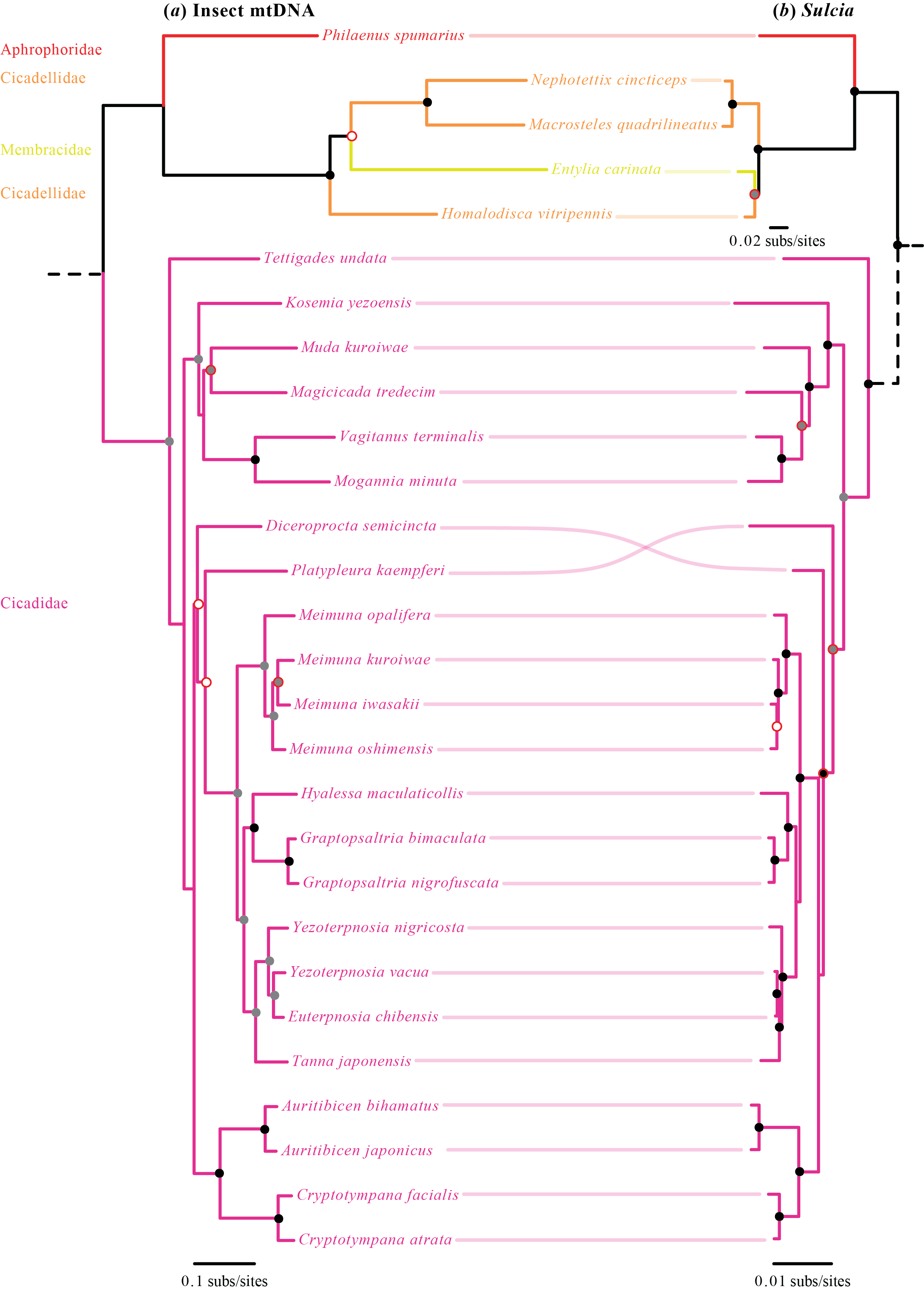

**Supplementary Fig. S8** Congruence between (*a*) phylogenetic tree of *Sulcia* hosts inferred using maximum likelihood (RAxML) from mitochondrial protein-coding genes, and (*b*) phylogenetic tree of *Sulcia* inferred using maximum likelihood from 120 protein-coding genes (3rd codon sites excluded from both data sets). Shaded circles at nodes indicate bootstrap values (black = 100%, grey = 85–99%). Nodes without black or grey circles have bootstrap values <85%. Red outlines on circles indicate disagreement between the phylogenies. Colours of branches indicate different *Sulcia* host families

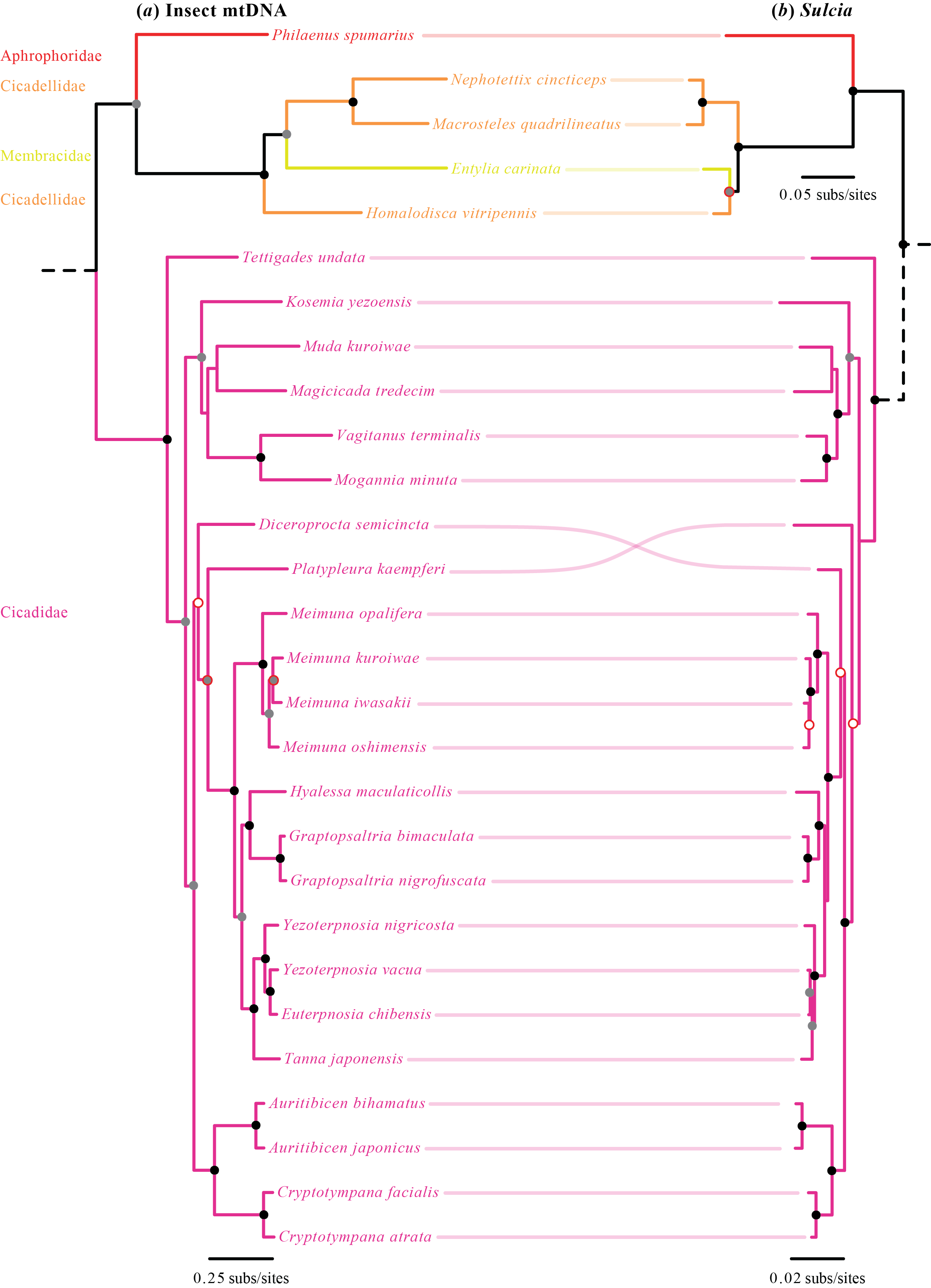

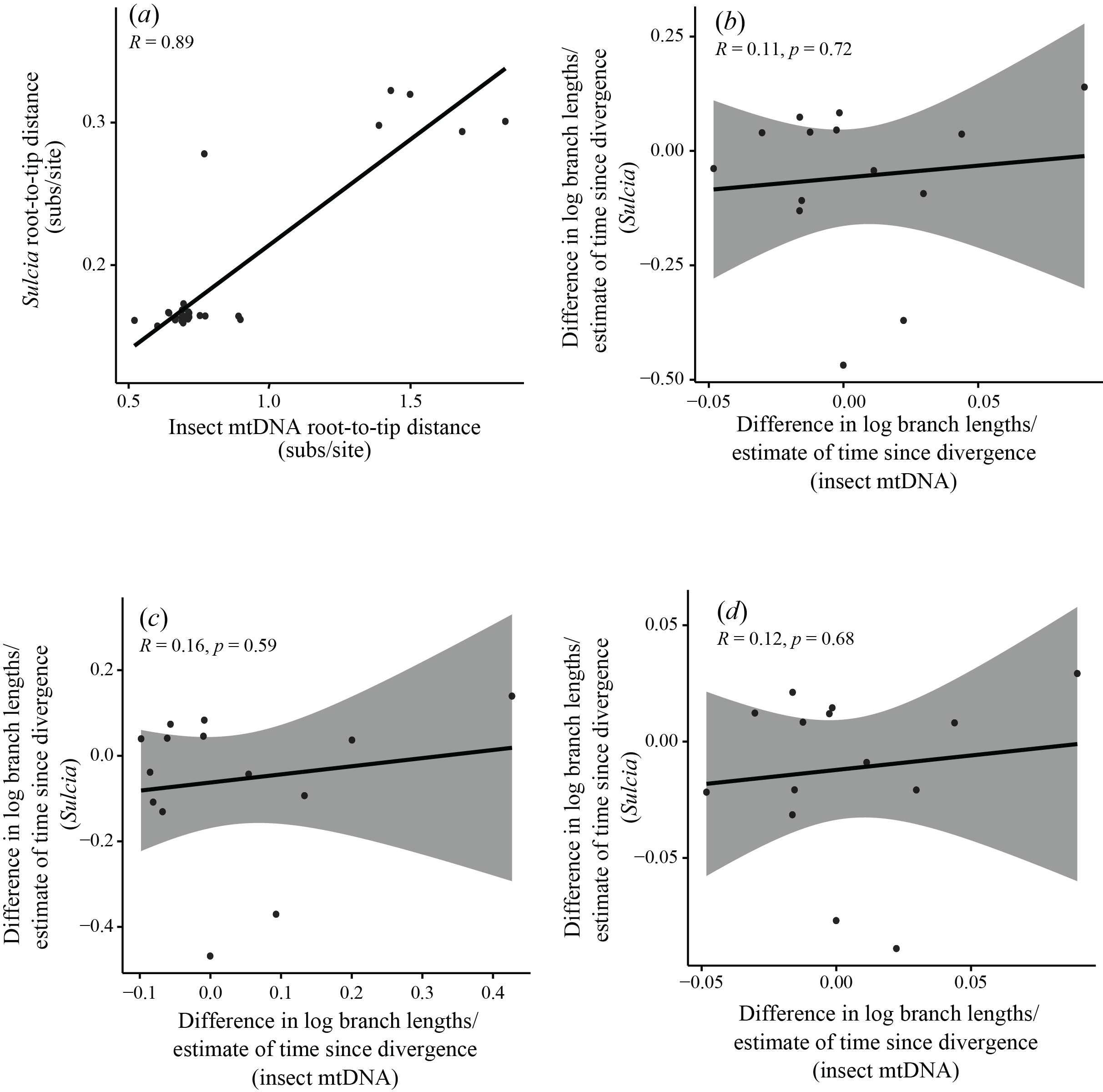

**Supplementary Fig. S9** Congruence between phylogenetic trees inferred using maximum likelihood (RAxML) from translated amino acid sequences of (*a*) mitochondrial protein-coding genes from *Sulcia* hosts, and (*b*) 120 protein-coding genes from *Sulcia*. Shaded circles at nodes indicate bootstrap values (black = 100%, grey = 85–99%). Nodes without black or grey circles have bootstrap values <85%. Red outlines on circles indicate disagreement between the phylogenies. Colours of branches indicate different *Sulcia* host families

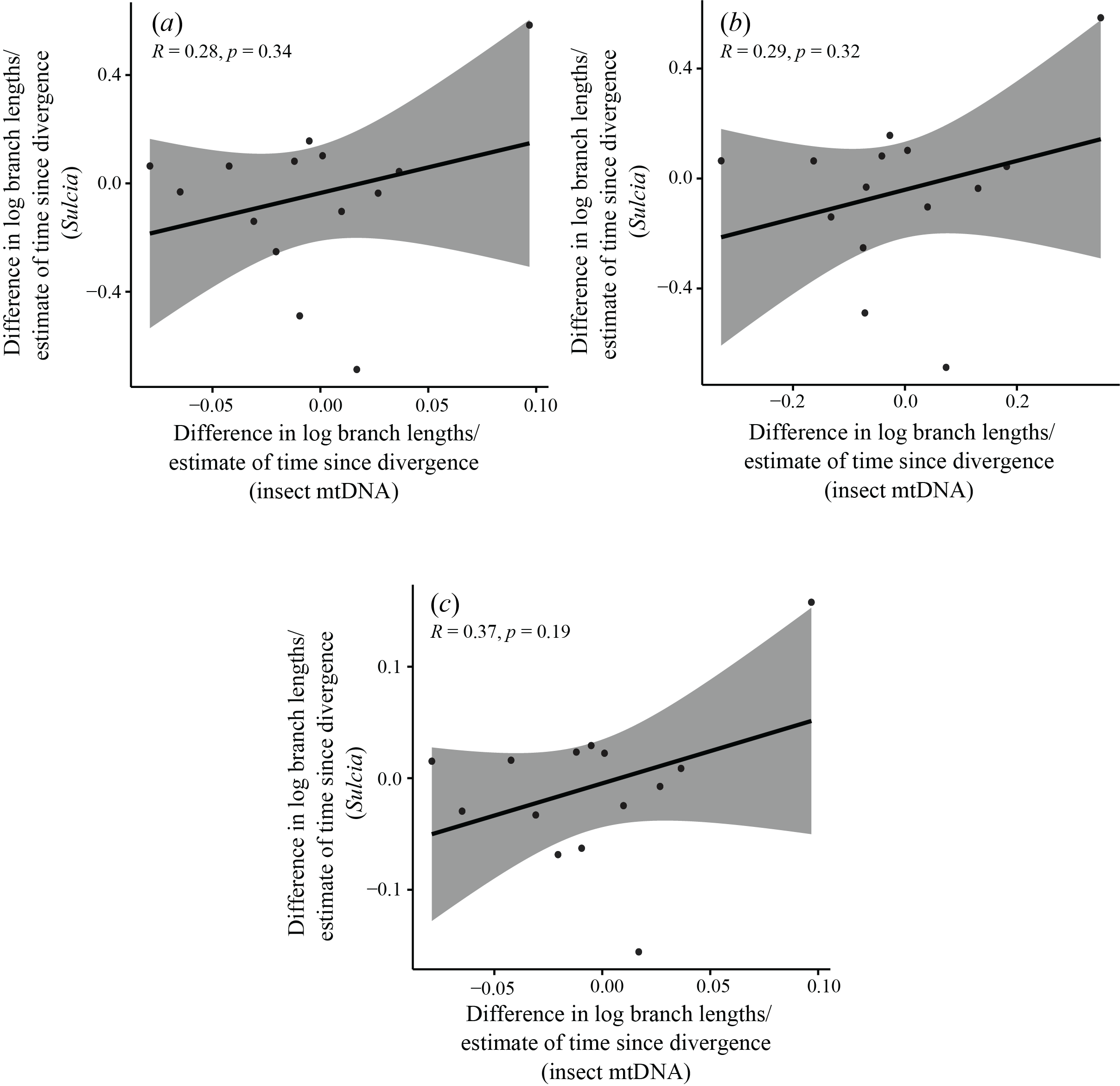

**Supplementary Fig. S10** Standardized tests for correlation of evolutionary rates between 14 independent pairs of *Sulcia* and host mitochondria, based on 120 symbiont genes and mitochondrial protein-coding host genes plus tRNAs, with 3rd codon sites excluded from host data sets. Three standardizations were carried out, each based on dividing log-transformed branch-length differences by the square root of an estimate of time since divergence for the pair. In the first standardization (*a*), time since divergence for host pairs was estimated as the average branch length of the host pair, divided by an assumed rate of 0.001 subs/site/million years, while for corresponding symbionts it was estimated as the average branch length of the symbiont pair, divided by the same assumed rate. In the second (*b*) and third (*c*) standardizations, times since divergence for both symbionts and hosts were based either on average branch lengths of host pairs only or symbiont pairs only

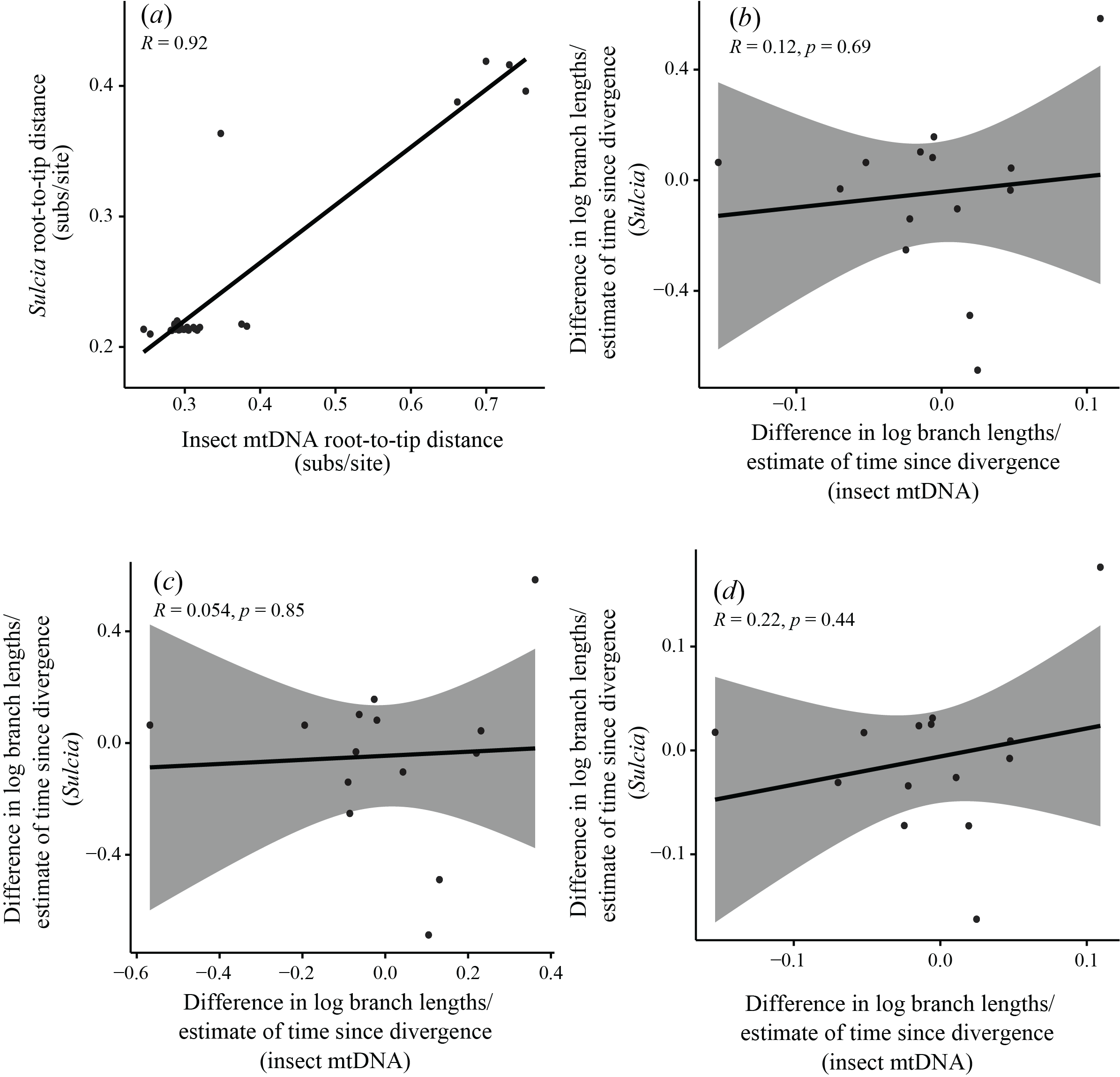

**Supplementary Fig. S11** Comparison of evolutionary rates of *Sulcia* symbionts and their hosts. (*a*) Correlation of root-to-tip distances in phylogenies of *Sulcia* and their hosts, inferred using maximum-likelihood analysis of protein-coding genes from each data set, with 3rd codon sites excluded from host data set. (*b–d*) Standardized tests for correlation of evolutionary rates between 14 independent pairs of *Sulcia* and host mitochondria. Three standardizations were carried out, each based on dividing log-transformed branch-length differences by the square root of an estimate of time since divergence for the pair. In the first (*b*), time since divergence for host pairs was estimated as the average branch length of the host pair, divided by an assumed rate of 0.001 subs/site/million years, while for corresponding symbionts it was estimated as the average branch length of the symbiont pair, divided by the same assumed rate. In the second (*c*) and third (*d*) standardizations, times since divergence for both symbionts and hosts were based either on average branch lengths of host pairs only or symbiont pairs only

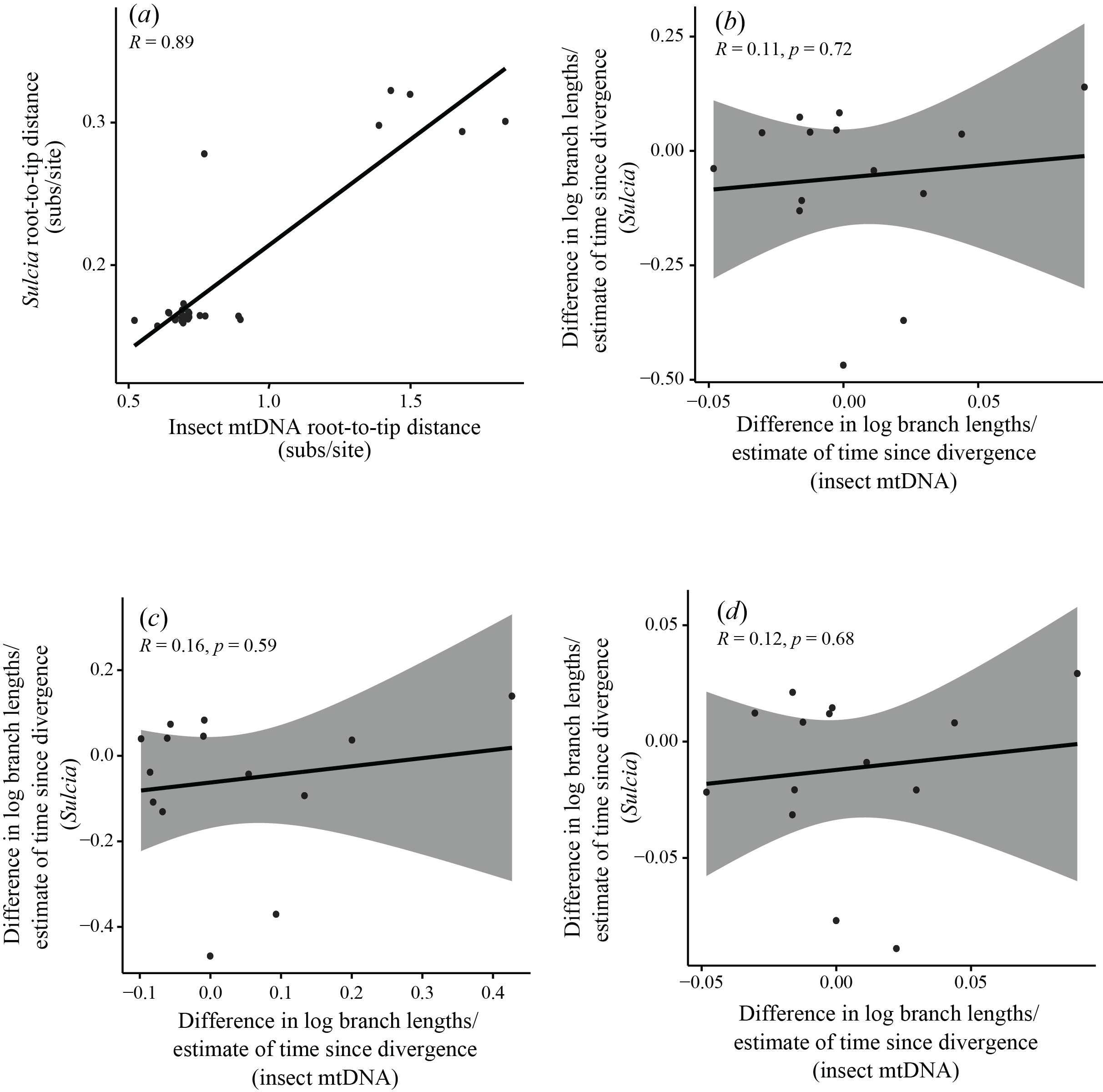

**Supplementary Fig. S12** Comparison of evolutionary rates of *Sulcia* symbionts and their hosts. (*a*) Correlation of root-to-tip distances in phylogenies of *Sulcia* and host, inferred using maximum-likelihood analysis of amino acid sequences translated from protein-coding genes. (*b–d*) Standardized tests for correlation of molecular evolutionary rates between 14 independent pairs of *Sulcia* and host mitochondria. Three standardizations were carried out, each based on dividing log-transformed branch-length differences by the square root of an estimate of time since divergence for the pair. In the first standardization (*b*), time since divergence for host pairs was estimated as the average branch length of the host pair, divided by an assumed rate of 0.001 subs/site/million years, while for corresponding symbionts it was estimated as the average branch length of the symbiont pair, divided by the same assumed rate. In the second (*c*) and third (*d*) standardizations, times since divergence for both symbionts and hosts were based either on average branch lengths of host pairs only or symbiont pairs only
