## Supplementary TableS1 for "Evolutionary rates are correlated between *Buchnera* endosymbionts and mitochondrial genomes of their aphid hosts"

**Table S1.** GenBank accession numbers for all samples used in this study

| Group | Species name | Symbiont | Host mtDNA |
| --- | --- | --- | --- |
| *Buchnera* | *Cinara tujafilina* | NC_015662 | KP722583 |
| *Buchnera* | *Anoecia oenotherae* | CP033012 | SRR7796603 |
| *Buchnera* | *Baizongia pistaciae* | NC_004545 | NC_035314 |
| *Buchnera* | *Schlechtendalia chinensis* | NZ_CP011299 | NC_032386 |
| *Buchnera* | *Melaphis rhois* | CP033004 | NC_036065 |
| *Buchnera* | *Thelaxes californica* | CP034852 | SRR7796612 |
| *Buchnera* | *Therioaphis trifolii* | CP032996 | SRR7796611 |
| *Buchnera* | *Stegophylla* sp. | CP032998 | SRR7796613 |
| *Buchnera* | *Tuberolachnus salignus* | NZ_LN890285 | KP722566 |
| *Buchnera* | *Muscaphis stroyani* | CP034861 | SRR7796591 |
| *Buchnera* | *Aphis craccivora* | CP034897 | KX447141 |
| *Buchnera* | *Aphis glycines* | NZ_CP009253 | KT889380 |
| *Buchnera* | *Aphis nasturtii* | CP034888 | SRR7796605 |
| *Buchnera* | *Aphis helianthi* | CP034894 | SRR7796607 |
| *Buchnera* | *Rhopalosiphum maidis* | CP032759 | MK3687781 |
| *Buchnera* | *Rhopalosiphum padi* | CP034858 | KT447631 |
| *Buchnera* | *Schizaphis graminum* | NC_004061 | NC_006158 |
| *Buchnera* | *Hyperomyzus lactucae* | CP034876 | MK2510631 |
| *Buchnera* | *Artemisaphis artemisicola* | CP034900 | SRR7796609 |
| *Buchnera* | *Acyrthosiphon pisum* | NC_002528 | NC_011594 |
| *Buchnera* | *Acyrthosiphon lactucae* | CP034891 | SRR7796608 |
| *Buchnera* | *Macrosiphum gaurae* | CP034867 | SRR7796596 |
| *Buchnera* | *Macrosiphum euphorbiae* | CP033006 | SRR7796595 |
| *Buchnera* | *Sitobion avenae* | CP034855 | NC_024683 |
| *Buchnera* | *Macrosiphoniella sanborni* | CP034864 | SRR7796594 |
| *Buchnera* | *Myzus persicae* | NZ_CP002701 | NC_029727 |
| *Buchnera* | *Brachycaudus cardui* | CP034879 | SRR7796601 |
| *Buchnera* | *Hyadaphis tataricae* | CP034873 | SRR7796597 |
| *Buchnera* | *Diuraphis noxia* | NZ_CP013259 | NC_022727 |
| *Buchnera* | *Lipaphis pseudobrassicae* | CP034870 | SRR7796598 |
| *Buchnera* | *Brevicoryne brassicae* | CP034882 | SRR7796604 |
| *Sulcia* | *Philaenus spumarius* | MPAX000000000 | AY630340 |
| *Sulcia* | *Nephotettix cincticeps* | CP016223 | KP749836 |
| *Sulcia* | *Macrosteles quadrilineatus* | CP006060 | NC_034781 |
| *Sulcia* | *Entylia carinata* | CP021172 | KX495488 |
| *Sulcia* | *Tettigades undata* | CP007234 | KJ193728 |
| *Sulcia* | *Kosemia yezoensis* | CP029015 | MG737723 |
| *Sulcia* | *Muda kuroiwae* | CP029017 | MG737729 |
| *Sulcia* | *Magicicada tredecim* | CP010828 | MG737744 |
| *Sulcia* | *Vagitanus terminalis* | CP029022 | MG737734 |
| *Sulcia* | *Mogannia minuta* | CP029016 | MG737728 |
| *Sulcia* | *Diceroprocta semicincta* | CP001605 | KM000131 |
| *Sulcia* | *Platypleura kaempferi* | CP029019 | MG737730 |
| *Sulcia* | *Meimuna opalifera* | CP029027 | MG737726 |
| *Sulcia* | *Meimuna kuroiwae* | CP029026 | MG737725 |
| *Sulcia* | *Meimuna iwasakii* | CP029025 | MG737724 |
| *Sulcia* | *Meimuna oshimensis* | CP029028 | MG737727 |
| *Sulcia* | *Hyalessa maculaticollis* | CP029014 | MG737722 |
| *Sulcia* | *Graptopsaltria bimaculata* | CP029012 | MG737720 |
| *Sulcia* | *Graptopsaltria nigrofuscata* | CP029013 | MG737721 |
| *Sulcia* | *Yezoterpnosia nigricosta* | CP029023 | MG737732 |
| *Sulcia* | *Yezoterpnosia vacua* | CP029024 | MG737733 |
| *Sulcia* | *Euterpnosia chibensis* | CP029011 | MG737719 |
| *Sulcia* | *Tanna japonensis* | CP029018 | MG737731 |
| *Sulcia* | *Auritibicen bihamatus* | CP029020 | MG737715 |
| *Sulcia* | *Auritibicen japonicus* | CP029021 | MG737716 |
| *Sulcia* | *Cryptotympana facialis* | CP029010 | MG737718 |
| *Sulcia* | *Cryptotympana atrata* | CP029009 | MG737717 |
